# Transitioning between preparatory and precisely sequenced neuronal activity in production of a skilled behavior

**DOI:** 10.1101/481960

**Authors:** Vamsi K. Daliparthi, Ryosuke O. Tachibana, Brenton G. Cooper, Richard H.R. Hahnloser, Satoshi Kojima, Samuel J. Sober, Todd F. Roberts

## Abstract

Precise neural sequences are associated with the production of well-learned skilled behaviors. Yet, how neural sequences arise in the brain remains unclear. In songbirds, premotor projection neurons in the cortical song nucleus HVC are necessary for producing learned song and exhibit precise sequential activity during singing. Using cell-type specific calcium imaging we identify populations of HVC premotor neurons associated with the beginning and ending of singing-related neural sequences. We discovered neurons that bookend singing-related sequences and neuronal populations that transition from sparse preparatory activity prior to song to precise neural sequences during singing. Recordings from downstream premotor neurons or the respiratory system suggest that pre-song activity may be involved in motor preparation to sing. These findings reveal population mechanisms associated with moving from non-vocal to vocal behavioral states and suggest that precise neural sequences begin and end as part of orchestrated activity across functionally diverse populations of cortical premotor neurons.

## INTRODUCTION

The sequential activation of neurons is implicated in a wide variety of behaviors, ranging from episodic memory encoding and sensory processing to the voluntary production of skilled motor behaviors^1-10^. Neural sequences develop through experience and have been described in several brain areas, including the motor cortex, hippocampus, cerebellum, and the basal ganglia^3,7,11-20^. Although computational models provide important insights into circuit architectures capable of sustaining sequenced activity^9,10,20-24^, our understanding of sequence initiation and termination is still limited.

The precise neural sequences associated with birdsong provide a useful biological model for examining this issue. Premotor projection neurons in the cortical vocal region HVC (HVC_RA_ neurons, see legend Figure 1 for anatomical abbreviations) exhibit precise sequential activity during song and current evidence suggests that this activity is acutely necessary for song production^1,25-28^. HVC_RA_ neurons are thought to be exclusively active during vocal production, yet ^~^50% of recorded HVC_RA_ neurons do not exhibit any activity during singing^1,6,27-29^, leaving the function of much of this circuit unresolved. To examine the neural mechanisms associated with the initiation and termination of singing we imaged from populations of HVC_RA_ neurons in freely singing birds.

**Figure 1.**
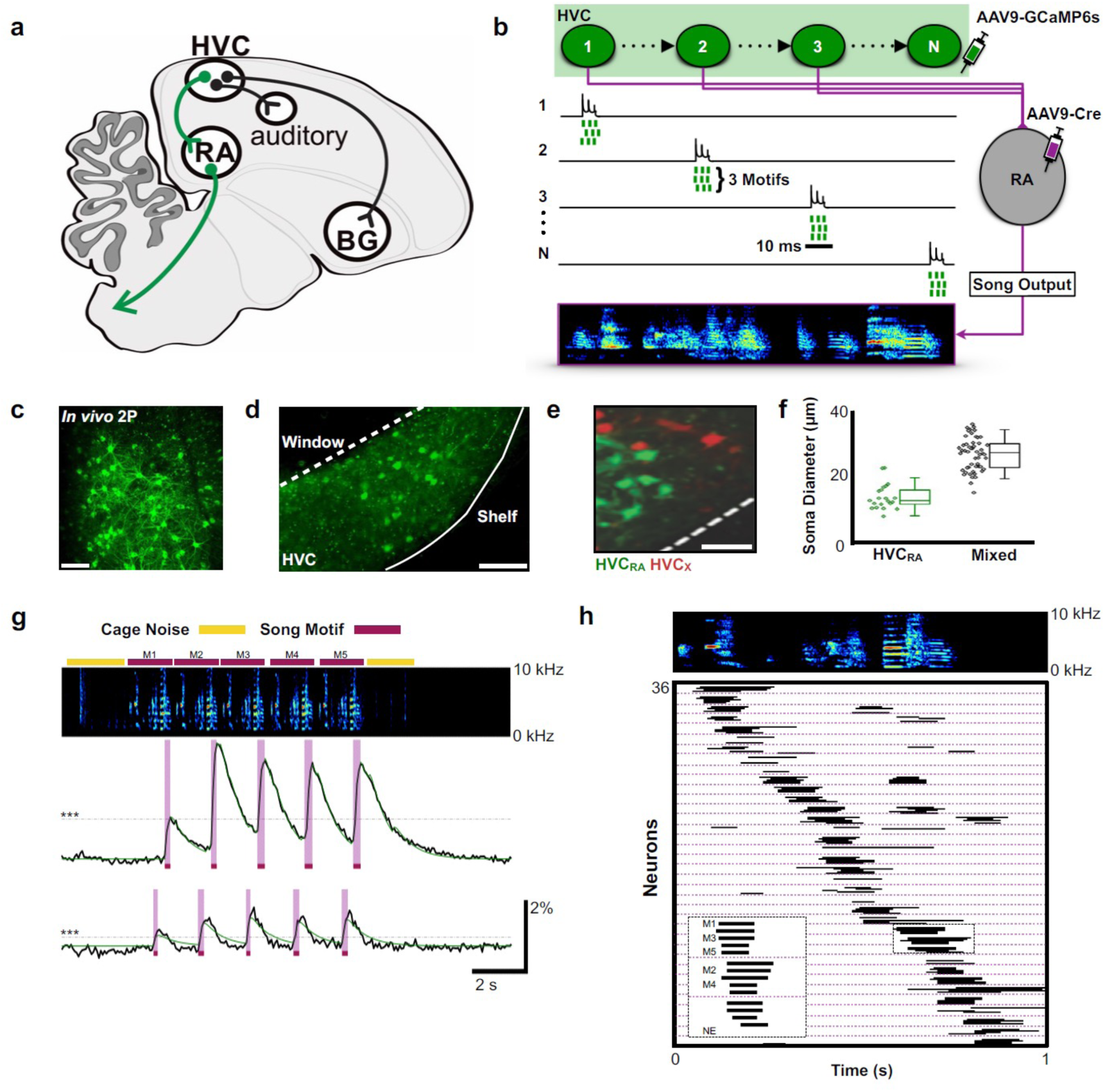
Population imaging of song-related HVC_RA_ sequences. **a)** Diagram showing 3 distinct projection neuron targets of the vocal premotor nucleus HVC. The projection neurons connecting HVC to the downstream motor nucleus RA (HVC_RA_ neurons) are shown in green. Basal ganglia (BG), robust nucleus of the arcopallium (RA), nucleus HVC of the nidopallium (HVC). **b)** Schematic showing HVC_RA_ neuron somata (green) and their outputs (magenta) to the downstream motor nucleus RA. AAV9-Flex-CAG-GCaMP6s was injected into HVC (green syringe) and AAV9-CAG-Cre was injected into RA (magenta syringe) to selectively label HVC_RA_ neurons. **c)** In vivo two-photon maximum density projection of retrogradely labeled HVC_RA_ neurons expressing GCaMP6s. Scale bar = 100 µm. **d)** A cross-section of HVC showing GCaMP6s-labeled HVC_RA_ neurons (green). The dashed line indicates where the cranial window was made over HVC. Note the lack of labeling in the region directly ventral of HVC, known as the HVC shelf. Scale bar = 100 µm. **e)** A sagittal section of HVC showing HVC_RA_ neurons (green) and retrogradely labeled HVC_X_ neurons (red). The dashed line indicates the border between HVC and HVC shelf. Scale bar = 50 µm. **f)** Whisker and scatter plots of soma diameters of GCaMP6s expressing cells show that retrogradely labeled HVC_RA_ neurons (green) have smaller diameters than neurons labelled using only direct viral injections into HVC (mixed population neurons, black). Boxes depict 25^th^ and 75^th^ percentiles, whiskers depict SD. HVC_RA_: N=21 neurons; mean diameter = 14.0 ± 3.8 µm (SD); Mixed: N=52 neurons; mean=26.9 ± 4.9 µm; *t* = -10.7, *p* = 2.0 × 10^-16^, two-sample *t* test. **g)** Example calcium traces from 2 HVC_RA_ neurons in a bird that sang 5 consecutive motifs. Shown are the background-subtracted traces (black) and the inferred calcium traces (green). The magenta overlays indicate the rise periods (intervals between onset and peak times) of the recorded calcium transients. The horizontal dashed line (gray) denotes 3 SD above baseline activity. The bars above the spectrogram denote cage noise associated with birds hopping or flapping their wings (yellow) or production of song motifs (red). **h)** Motif-related activity of 36 HVC_RA_ neurons across 5 motifs. Each row shows activity of a neuron from 1 trial. The dashed magenta lines separate different neurons. Empty spaces indicate trials wherein neurons were not active (no event, NE). The inset shows a zoom-in of activity from 3 separate HVC_RA_ neurons.

Motor planning and preparation activity are associated with accurate production of volitional motor movements^8,30^ but are poorly described in the context of precise neural sequences or in the production of song. Here we show that ^~^50% of HVC_RA_ neurons are active during periods associated with preparation to sing and recovery from singing. One population is only active immediately preceding and following song production, but not during either singing or non-vocal behaviors. A second population of neurons exhibits ramping activity before and after singing and can also participate in precise neural sequences during song performance. Recordings from downstream neurons in the motor cortical nucleus RA reveal neural activity prior to song initiation and during song termination. The control of respiratory timing is essential for song^31^, and our measurements of respiratory activity suggest that pre-singing activity in HVC_RA_ neurons functions to coordinate changes in respiration necessary for song initiation. From these findings, we argue that subpopulations in HVC encode the neural antecedents of song that drive recurrent pathways through the brainstem to prepare the motor periphery for song production.

## RESULTS

### Activity Sequences in Populations of HVC_RA_ Neurons

We used miniscope calcium imaging to examine the activity of populations of HVC_RA_ neurons in singing zebra finches^32,33^. A total of 223 HVC_RA_ neurons were imaged during production of 1,298 song syllables from 6 birds (30 song phrases across 18 imaging trials, **EDTable 1**). To selectively target HVC_RA_ neurons, we combined retrograde viral expression of cre recombinase from injections into RA with viral expression of cre-dependent GCaMP6s from injections into HVC (**Figure 1a-b and legend, see Methods**)^33^. We confirmed the identity of imaged neurons using conventional retrograde tracing, anatomical measures of neuronal features, and post-hoc histological verification. We found that this approach exclusively and uniformly labeled populations of HVC_RA_ neurons (**Figure 1c-f**).

To elicit courtship singing, we presented male birds with a female and imaged HVC_RA_ neurons during song performance (**EDVideo 1**). Given the slow decay times of calcium signals relative to singing behavior, we defined neuronal activity by the rise times of calcium events that were >3 standard deviations (SD) above baseline (**Figure 1g,** average rise time: 0.112 ± 0.047 s SD, **see Methods**). The activity of individual HVC_RA_ neurons was time-locked to a moment in the birds’ song, with different neurons active at different moments in the song motif (**Figure 1g-h**; onset jitter = 55.0 ± 60.9 ms, populations imaged at 30 frames per second). We found that the sequential activity of HVC_RA_ neurons roughly coded for all moments in the song motif (**Figure 1h**). These results provide the first glimpse of activity across populations of identified HVC_RA_ neurons during singing and support the idea that sparse and precise neuronal sequences underlie the sequential structure of birdsong^1^,^6^,^27^,^34^,^35^.

### Peri-Song Activity in Populations of HVC_RA_ Neurons

The sequence of syllables in zebra finch song is stereotyped and unfolds in less than a second. Like other rapid and precise motor movements, song may benefit from motor planning and preparatory activity; however, HVC_RA_ neurons have been hypothesized to exclusively represent temporal sequences for songs and calls, and to remain quiescent at all other times^1,5,6,28,29,34^. To scrutinize this prediction, we examined the activity of HVC_RA_ neurons prior to song onset, in between song bouts, and immediately after singing (**Figure 2a-c, EDFig. 1,** song onset defined as the beginning of a song-phrase, including introductory notes; see **Methods** for detailed definitions of phrases, song, and peri-song behavior). For any given song phrase, we observed significant activity surrounding singing behavior, refuting the hypothesis that HVC_RA_ neuronal activity is restricted to periods of active song production. 30 out of 30 song phrases from 6 birds exhibited ‘peri-song’ activity, defined as the 5 second intervals before and after singing, plus silent gaps within song phrases. Slightly fewer HVC_RA_ neurons were active during peri-song intervals (54.3% of neurons) than during singing (59.9%), (*t* = -1.1, *p* = 0.29 two-sample *t* test; **Figure 2b)**. Of the neurons that were active during peri-song intervals, 40.6% were active prior to song onset, 17.8% were active following song offset, and 41.6% were active before and after song (N=197 neurons, 6 birds, **EDFig. 2**). When we examined the timing of peri-song events, we found no correlation in the timing of pre-song and post-song event onsets in neurons that were sparsely both before and after song (**EDFig. 3**). Although neuronal populations displayed considerably more calcium events during song (*t* = 5.635, *p* = 5.65 x10^-7^ two-sample *t*, normalized song event rate = 24.09 events/s ± 15.95 SD, normalized peri-song event rate = 6.97 events/s ± 3.72 SD), a substantial fraction of all recorded calcium events occurred within the 5 second intervals before or after song (997/2366 or 32.5% of all calcium events).

**Figure 2.**
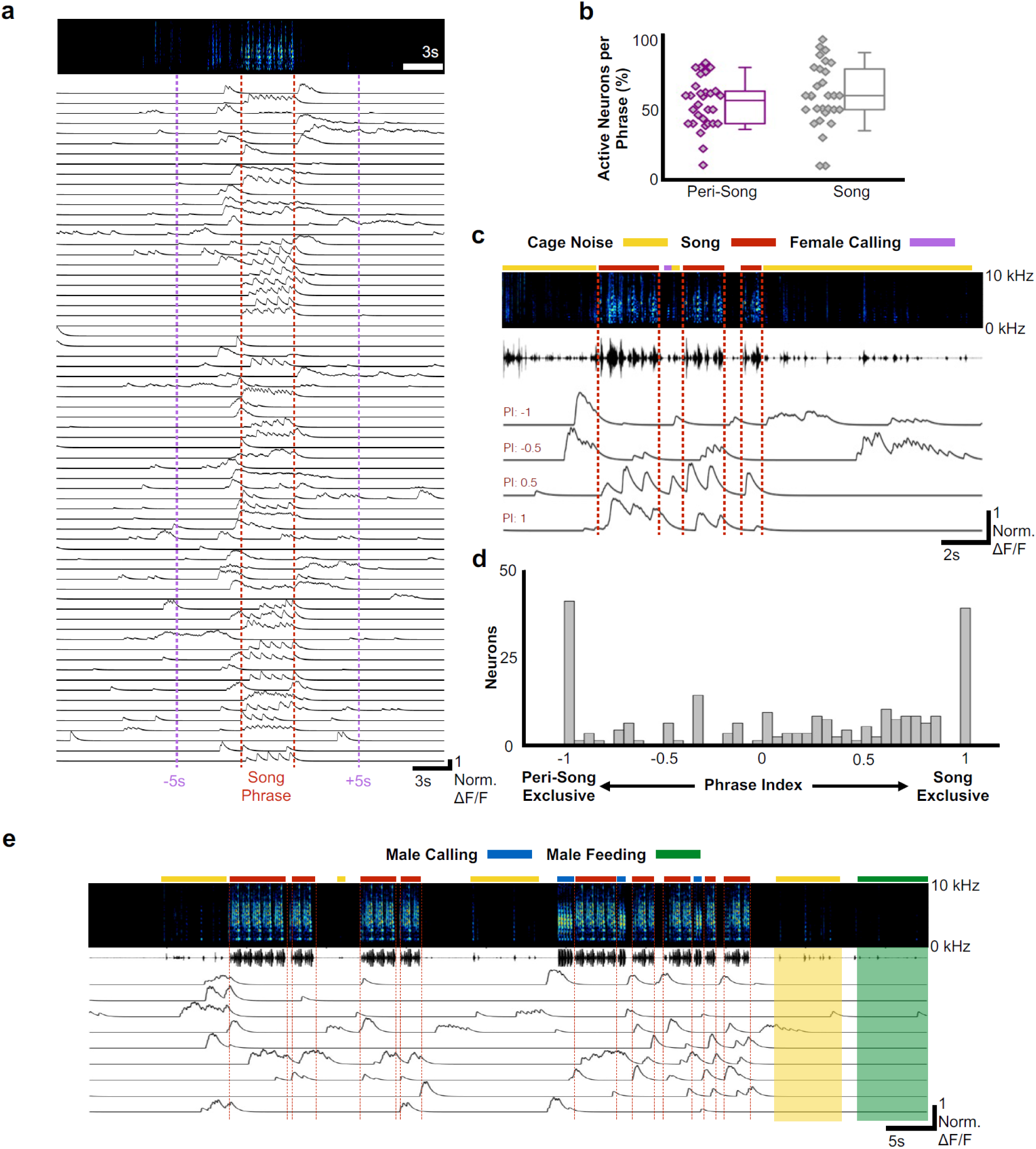
Most HVC_RA_ neurons exhibit peri-song activity. **a)** Normalized calcium transients from 67 simultaneously recorded HVC_RA_ neurons during production of a song phrase. The red dashed lines delimit 5 consecutive motifs. **b)** Percentages of active neurons during peri-song (54.3 ± 17.8 %, SD, purple) and song (59.9 ± 22.5 %, gray) are similar (30 phrases, *t* = -1.1, *p* = 0.29 paired two-sample *t* test). Box plots show the median, 25^th^ and 75^th^ percentiles with whiskers showing ±1.5 IQR. **c)** Sample neurons with diverse phrase indices ranging from -1 to 1 and their corresponding calcium traces during 6 motifs over 3 bouts. Dashed lines indicate bout onsets and offsets. Bars above spectrogram indicate the presence of cage noise related to hopping and wing flapping (yellow) or female calling (FC, blue). **d)** Histogram of phrase indices for all 223 neurons from 6 birds. **e)** Undirected song from a different male showing periods of cage noise or hopping behavior (yellow) and feeding behavior (green). Blue boxes indicate the male calling. Red dashed lines indicate onsets and offsets of song bouts.

**Figure 3.**
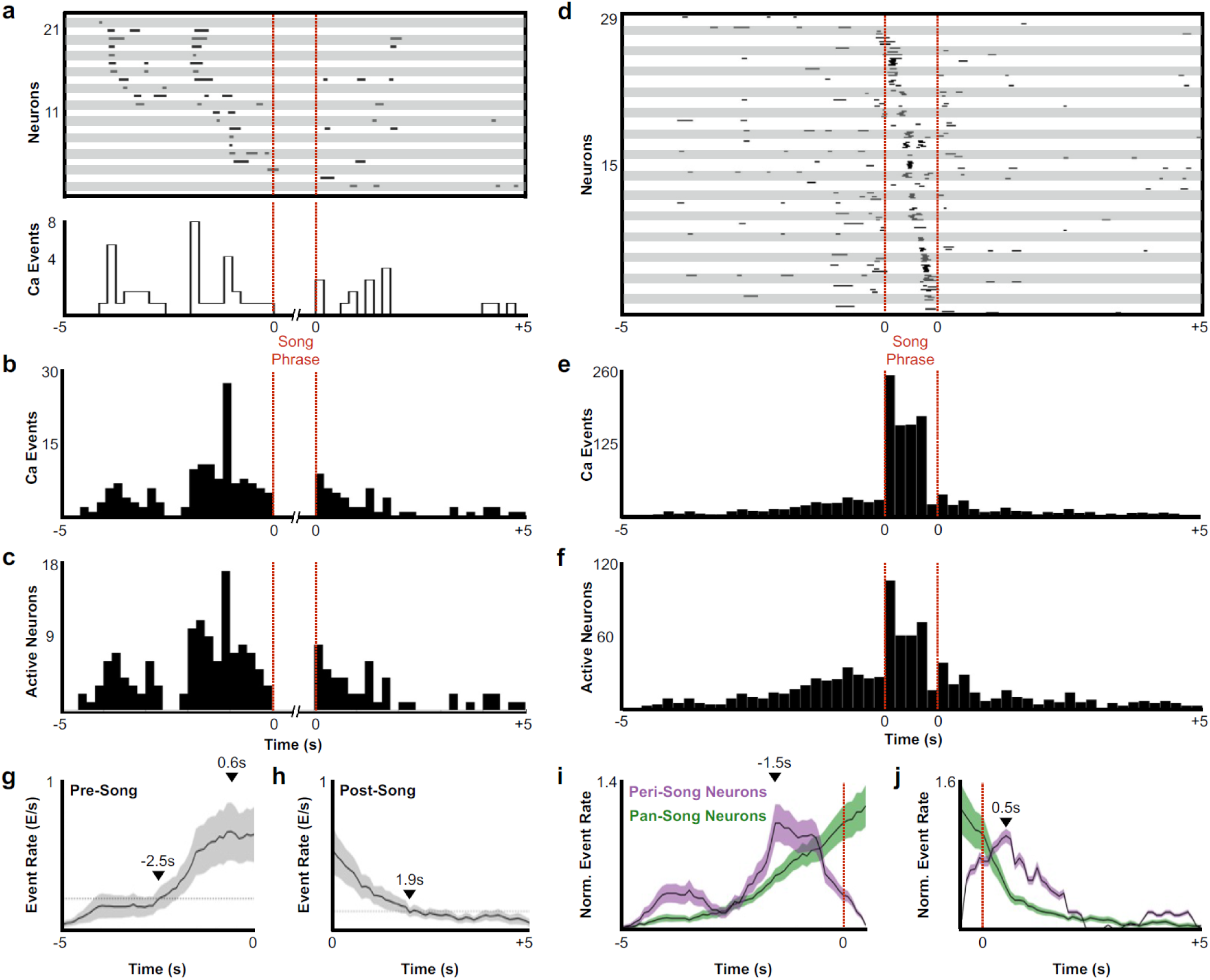
Description of peri-song and pan-song neuron activity. **a)** 21 peri-song neurons from one bird singing 3 bouts containing 6 motifs (1st bout: 3 motifs; 2nd bout: 2 motifs; 3rd bout: 1 motif, the dashed red lines indicate the onset and offset of the three bouts). Each row shows song-aligned calcium events (CEs N=54 CEs; average rise time = 0.18 ± 0.09 s SD). The shaded horizontal bars separate different neurons. One CE is seen to overlap with the beginning of the song phrase. The onset time for this event is 170 ms before song, but the rise time is slow and extends to 100 ms after song onset. Below the CE raster plot is a peri-event histogram with the event rate in 200 ms bins shown for the trial above. **b)** Song-aligned CEs in peri-song neurons 5 s before and after phrase onset (41 neurons, 190 CEs). The activity rate peaks ^~^1.2 s before phrase onset. **c)** The number of active peri-song neurons in 200 ms bins before and after phrase onset (41 neurons, 169 CEs). **d)** Song-aligned activity of pan-song neurons (same trial as shown in panel **a**, N=29 neurons, 253 CEs). **e)** Peri-event histogram of pan-song neurons (143 neurons, 1,333 CEs). **f)** The number of active pan-song neurons in 200 ms bins (143 neurons, 853 CEs). **g)** Pre-phrase event rate for all pan-song neurons. The event rate was calculated by counting event onsets in 100 ms bins and then smoothed with a 1 second moving window. 28 trials are shown from 5 birds, the black line indicates the average event rate. The black triangles mark the peak event rate occurring 0.6 s before song onset and 2.5 s when the event rate reaches 3 SD above the baseline event rate, respectively. Baseline event rate was determined by measuring the average event rate from -5 to -4 seconds before song onset. Shaded region indicates standard deviation. **h)** Post-phrase event rate for all neurons. 27 trials are shown from 5 birds. The black triangles mark when the event rate reaches 3 SD above the baseline event rate. Baseline event rate was determined by measuring the average event rate during -5 to -4 seconds before song onset. **i)** Pre-phrase event rate for peri-song and pan-song neurons calculated as calcium events in moving 1 s windows. The black line indicates average event rate. The black triangle indicates peak event rate occurring 1.5 s before phrase onset. **j)** Same as **i**, but post-phrase event rates for peri-song and pan-song neurons. The black triangle indicates peak event rate occurring 0.5 s after phrase offset.

To better characterize these newly discovered activity profiles, we indexed the song and peri-song activity of all HVC_RA_ neurons throughout a day of singing (phrase index: range -1 to +1, with neurons exclusively active outside of singing scoring -1 and neurons active only during singing +1, **Figure 2c-d, see Methods**). We found that HVC_RA_ phrase indices were not uniformly distributed (χ^2^ (7, N = 223) = 46.3, *p* = 7.6 × 10^-8^, Chi-square goodness of fit test), with a significant fraction (36%) falling at the extremes of this scale (**Figure 2d**), and that neurons with different phrase indices were anatomically intermingled throughout HVC (**EDFig. 4**). Most neurons (64.1%) displayed sparse heterogenous activity during peri-song periods and transitioned to temporally precise activity during singing (referred to here as ‘**pan-song neurons**’; **EDFig. 5**). At the extremes of the phrase index scale (phrase indices -1 or +1), we found that 18.4% of neurons were exclusively active during peri-song intervals (phrase index = -1, referred to here as ‘**peri-song neurons**’), while 17.5% participated exclusively in neural sequences during singing (phrase index = +1, referred to here as ‘**song neurons**’).

**Figure 4.**
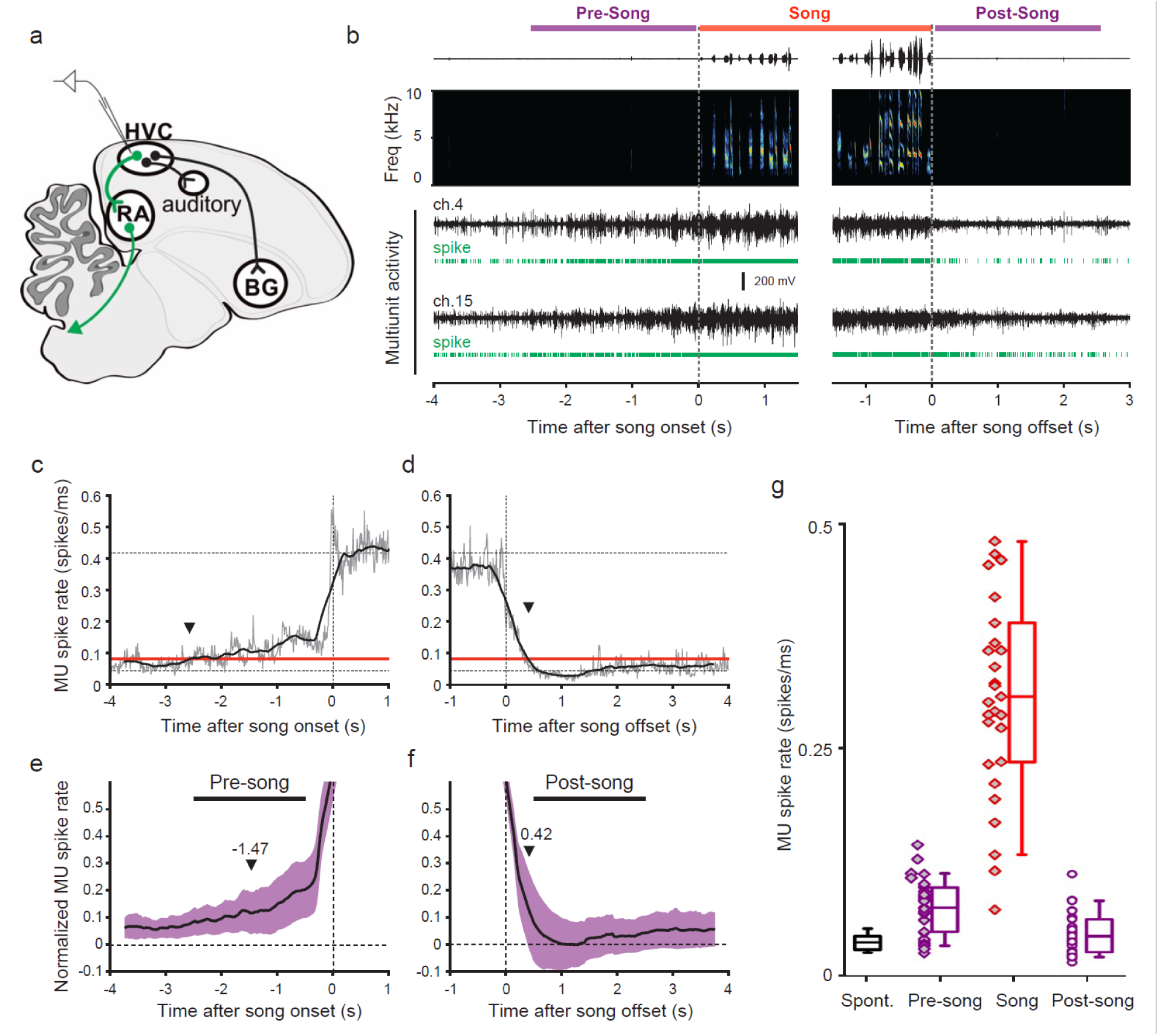
Pre-song and post-song firing in HVC of Bengalese finches. **a)** Schematic of recording site. **b)** An example of song initiation and termination (dashed lines indicate phrase onset and offset, pre and post-song period marked in purple and song marked in red) and simultaneously recorded HVC multiunit activity on two electrodes (channels #4 and #15, bird p15o56). Green raster plots represent detected spikes on the two electrodes. **c,d)** Phrase onset (c) and offset (d) related average multiunit activity obtained from an example electrode channel (vertical dashed lines indicate phrase onset and offset, respectively). The spike rate was averaged across multiple song onsets or offsets and was first calculated in 10 ms bins (gray thin line) and then smoothed with a 500 ms window (bold line). Upper and lower horizontal dotted lines show mean spike rates during singing and baseline, respectively. Arrowheads indicate onset (c) and offset (d) timings of spike rate, as assessed by crossing of a pre-defined threshold (red line). **e,f)** Normalized multiunit activity related to phrase onset (e) and offset (f). Before averaging, the spike rate trace of each electrode channel was normalized such that 0 corresponds to the mean rate during baseline and 1.0 to the mean rate during singing (see Method). The bold line shows an average across all electrodes and birds (n = 29 channels). Purple area indicates ± one SD. Arrowheads show mean onset (e) and offset (f) timing of pre-song and post-song activity, respectively. **g)** Mean multiunit spike rates during spontaneous (black), pre-song (purple with diamonds), song (red), and post-song (purple with circles) periods. Pre-song and post-song periods are indicated by the horizontal bars in panels e and f. Box plots show the median, 25^th^ and 75^th^ percentiles with whiskers showing ±1.5 IQR.

**Figure 5.**
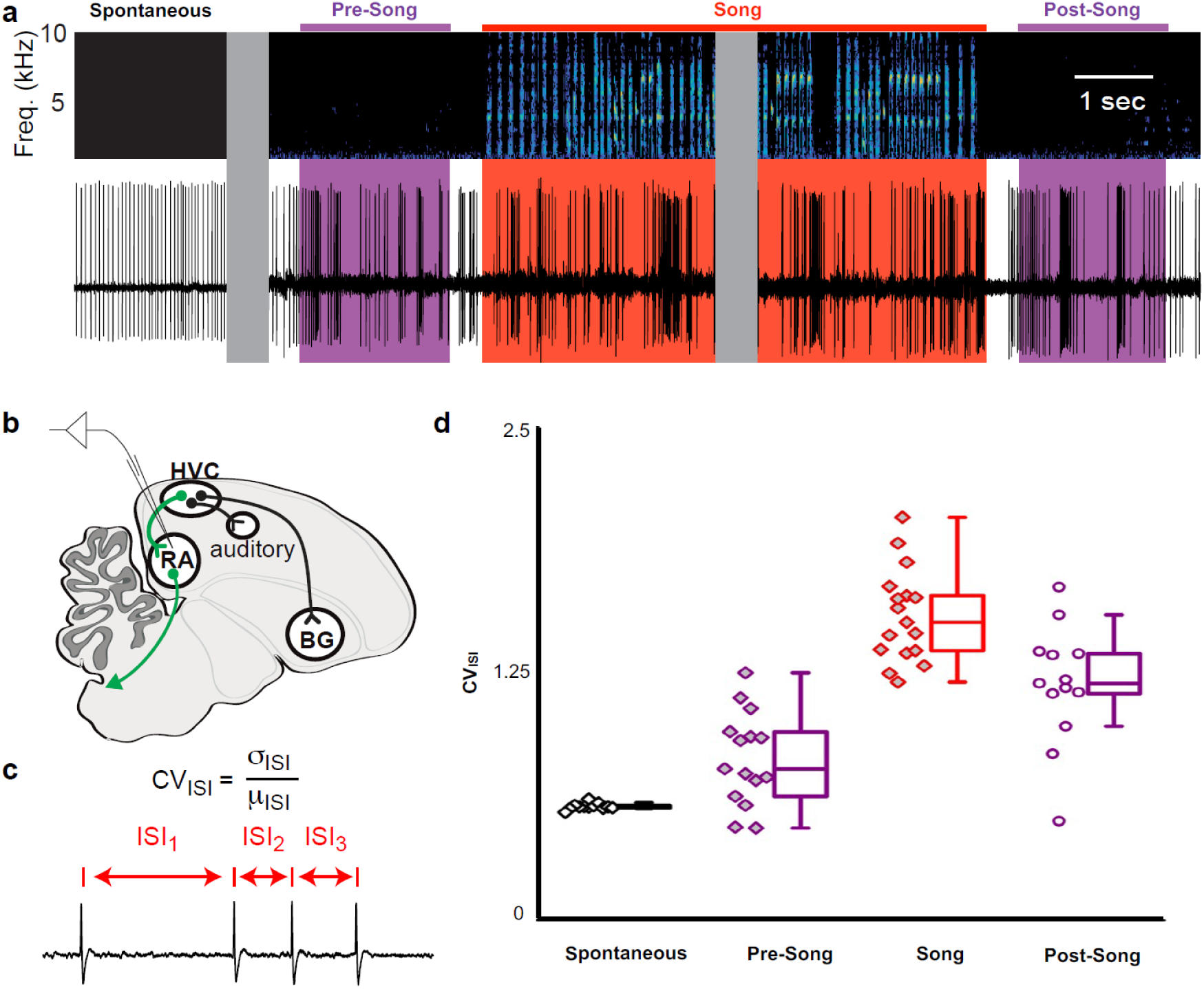
**a)** Example extracellular recording from a single RA neuron. Colored areas highlight four epochs (spontaneous, pre-song (purple), song (red), and post-song (purple)) relative to the beginning and ending of a song phrase (see main text). Grey areas indicate discontinuities in time (pauses between “spontaneous” epoch and song initiation and within the middle portion of the song bout). **b)** Schematic of recording site. **c)** We quantified inter-spike-intervals (ISIs) and computed the coefficient of variation (CV) in each epoch. **d)** We found significantly higher ISI variability in the pre-song epoch (purple with diamonds) compared to spontaneous (*p*<0.005, two-sided K-S test). Box plots show the median, 25^th^ and 75^th^ percentiles with whiskers showing ±1.5 IQR.

These results provide a starkly different view of HVC_RA_’s potential contribution to behavior, revealing that a substantial portion of this network can be active outside of the precise neuronal sequences associated with song. One interpretation is that HVC_RA_ neurons may play a role in motor planning and in preparation to sing. That more than half of all HVC_RA_ neurons can be active during peri-song intervals also raises the prospect that precise neural sequences emerge as part of changing network dynamics across subpopulations of HVC_RA_ neurons. Lastly, these results lend insight into why approximately half of HVC_RA_ neurons recorded using electrophysiological methods appear to be inactive during song. Nonetheless, this result also raises questions as to why previous studies have not identified peri-song activity. First, previous calcium imaging experiments have not restricted GCaMP expression to HVC_RA_ neurons, but rather relied on either non-selective labeling of neuronal populations in HVC^4,34,36,37^ or have been restricted to imaging small populations of other classes of HVC neurons^38^. Second, electrophysiological studies of identified HVC_RA_ neurons have been mostly confined to recording one neuron at a time and these experiments have focused on understanding coding during song production, often using short (^~^500 ms) buffering windows triggered by singing behavior. Therefore, sparse heterogenous activity occurring seconds before or after song could be simply overlooked or could appear irrelevant unless viewed through the lens of population dynamics. Consistent with this idea, previous multi-unit recordings in zebra finches and mockingbirds, which are dominated by activity of interneurons or neurons projecting to the basal ganglia, have identified ‘anticipatory’ activity in HVC hundreds of milliseconds prior to song onset, but the role of this activity and whether HVC_RA_ neurons are active prior to song onset have not been examined^39-41^.

Another possibility is that peri-song activity is unrelated to singing and merely reflects low levels of spontaneous activity intrinsic to HVC_RA_ neurons. This was not the case, however, as HVC_RA_ neurons were largely inactive outside of pan-song intervals and were significantly more active during the peri-song periods than baseline (baseline calculated from periods ≥10 s removed from periods of singing or calling, *p* = 8.4 × 10^-5^, Chi-square = 18.78 Friedman test, baseline = 12.9% ±5.7 SD of fluorescence values normalized to song, pre-song = 26.9% ±14.7, and post-song = 30.2% ±10.7). In addition, we examined the amplitudes of peri-song calcium events and found that they were larger than events occurring during song (*t* = 3.2769, *p* = 0.0012, two-tailed *t* test, 279 fluorescence peaks measured from neurons active during both peri-song and song, pan-song neurons with phrase indices between -0.18 to 0.18). We also asked whether peri-song activity might relate to factors other than singing. We examined trials in which birds did not sing to female birds but found no HVC_RA_ neuron that responded solely to presentation of the female or during non-song related movements of the head, beak, or throat, such as during eating, grooming, and seed-shelling (**Figure 2e, EDFig. 6)**. Indeed, our populations of HVC_RA_ neurons only became substantially active prior to singing or calling. Moreover, we found peri-song activity in the moments before and after undirected song bouts produced in isolation (**EDFig. 6 - 7**), suggesting that peri-song activity is unlikely to be solely associated with extraneous courtship behaviors such as courtship dance.

**Figure 6.**
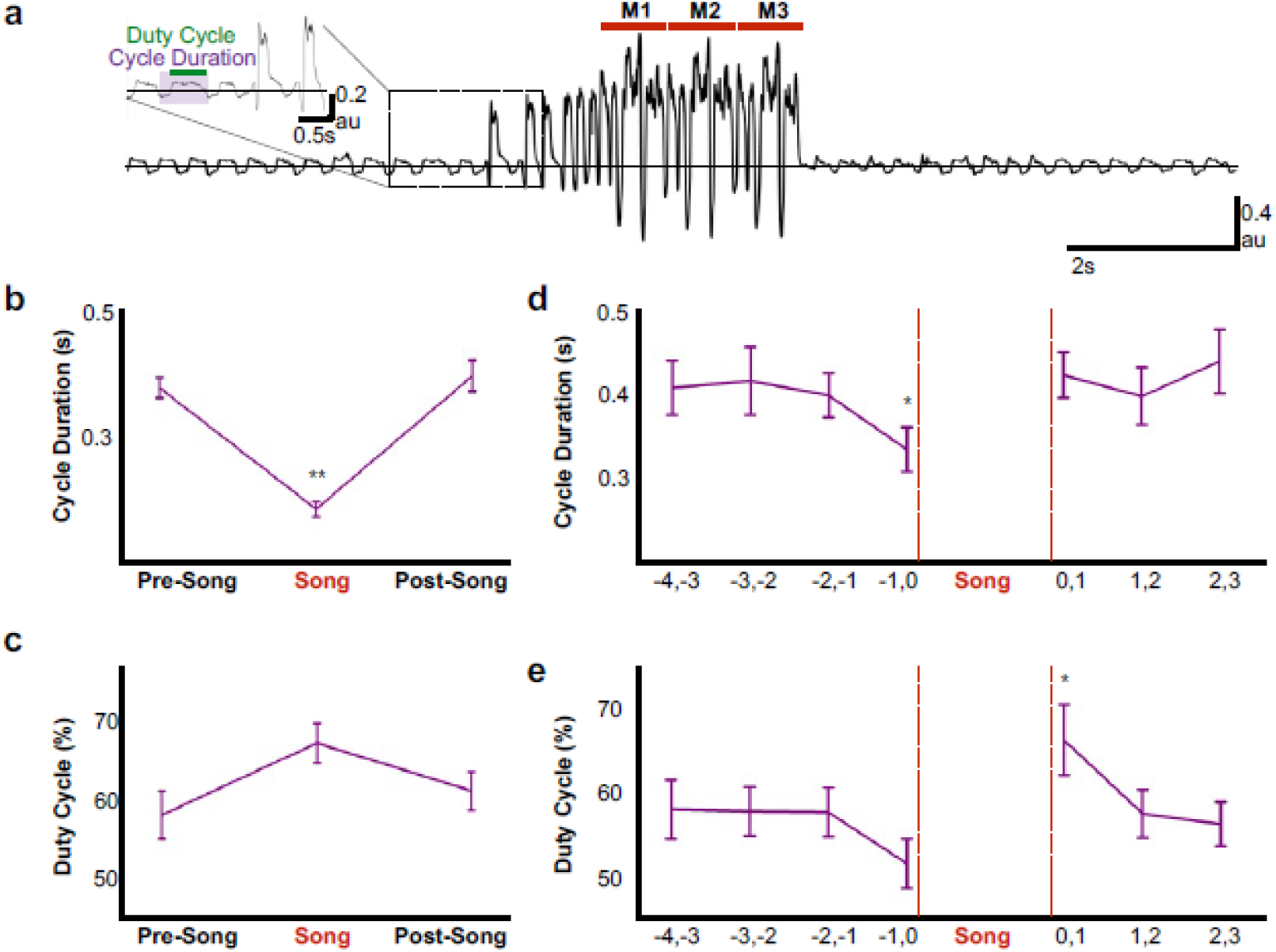
Air sac pressure recording in zebra finches. **a)** Waveform of pressure changes during non-singing and singing periods. Waveforms above the horizontal line (suprambient pressurization) indicate expiration and below the line (subatmospheric pressurization) indicate inhalation. Inset illustrates measurements for respiratory cycle duration and duty cycle (% time in expiration). **b)** Respiratory cycle duration and **c)** duty cycle of expiratory phase before (Pre), during (Song), and after (Post) song production (N = 6 birds). **d)** Plots of average respiratory cycle durations and **e)** duty cycles during pre-song and post song periods (N = 6 birds). Longer duty cycles correspond to increased periods of expiration.

We next examined the possibility that HVC_RA_ neurons play a role in motor planning or preparation as birds prepare to sing. We found pre-song activity in 28/30 song phrases analyzed. In the two instances when we did not detect any pre-song activity, less than 6 HVC_RA_ neurons were active within our imaging window during singing, indicating that the lack of activity was likely the result of under sampling from the population. Electrophysiological recordings in young zebra finches have identified HVC neurons that mark the onset of song bouts^5^, ‘bout neurons’ that burst immediately prior to vocalizations. The vast majority of pre-song activity we describe occurs hundreds of milliseconds to seconds prior to vocalizations, suggesting a role in planning or preparation to vocalize (96.2% of calcium events occurred more than 100 ms prior to vocal onset and 65.6% occurred more than 1 second prior to vocal onset). Zebra finches often sing a variable number of introductory notes prior to the first song motif of a song bout. Pre-song activity (prior to introductory notes) could be related to the number of introductory notes to be sung, but we found no correlation between pre-song event rates and the number of introductory notes (**EDFig. 8**). In 27/28 song phrases we found increases in population activity greater than 3 SDs above baseline predicted song onset within the following 4-5 seconds (2.44 s ±1.0 s SD, 28 song phrases from 5 birds). Calcium activity reached two-thirds of the maximum pre-song activity only prior to song onset or prior to short vocalizations (**EDFig. 9**). Together, these results indicate that HVC_RA_ neuron activity is predictive of the voluntary production of courtship song and suggests a role for this network in motor planning and in preparation to sing.

### Peri-Song and Pan-Song Neurons

A substantial fraction of all imaged neurons (41/223 neurons) were active exclusively before or after singing (**Figure 2d**, neurons with a phrase index of -1). These peri-song neurons exhibited sparse heterogeneous activity before song phrases and were occasionally active in the silent intervals between song motifs (**Figure 3a-c, and EDFig. 10**). Both the number of active peri-song neurons and the density of calcium events increased 1-3 s prior to song onset, with the event rate peaking 1.5 s prior to singing (**Figure 3b-c, i**) and then declining sharply in the last second before song onset (**Figure 3i**). Most of the neurons we imaged, pan-song neurons (143/223 neurons), exhibited sparse, heterogeneous activity before and/or after song and exhibited time-locked sequences during singing (**Figure 3d-f**). Pan-song neurons exhibited substantial increases in their activity in the last ^~^2 s prior to song onset. Their activity continued to increase as the activity of peri-song neurons began to ramp-off prior to song onset (**Figure 3g,I, and EDFig. 11**). Differences in pre-song activity profiles between peri-song and pan-song neurons (Kolmogorov-Smirnov, K-S test, *k* = 0.26, *p* = 0.056) may support temporally coordinated network transitions as birds prepare to sing, suggesting that neural sequences for song could emerge as part of changing network dynamics in HVC.

During production of song motifs, pan-song neurons exhibited sequential bursts of activity that roughly coded for all moments in the bird’s song, similar to sequencing previously described in song neurons. We found that pan-song and song neurons had a similar probability of being active with each motif (song neurons probability of at least one calcium event per motif P(motif) = 0.72 ± 0.32, pan-song neurons P(motif) = 0.66 ± 0.29, K-S test, *p* = 0.11; **EDFig. 12**); however, we also noted that the probability of being active was lower than in electrophysiological recordings^1,28^. This likely reflects limitations in event detection using single-photon calcium imaging. To better understand this, we calculated signal to noise ratios (SNRs, signal defined as peak fluorescence of calcium events during song) between pan-song neurons and song neurons during singing (SNR song neurons = 913.8 ±405.4 (7 neurons), pan-song neurons = 638.3±248.2 SEM, n=36 neurons). We found no difference in SNR between these neurons (two-sample t test, *p* = 0.6; SNR calculated from 156 calcium events (28 from song neurons and 128 from pan-song neurons), suggesting that although we are underestimating activity during singing, these limitations are unlikely to obscure differences between pan-song and song neurons.

In addition to preceding song onset, neurons also marked the end of song phrases. Within 5 seconds after song offset, peri-song neurons exhibited a sharp increase in activity followed by a gradual ramp-off (**Figure 3a-c, h, j, and EDFig. 11**) whereas pan-song neurons only exhibited a ramp-off (**Figure 3d-f, h, j**). Although the distribution of post-song activity differed between peri-song and pan-song neurons (K-S test, *k* = 0.2692, *p* = 0.0373), both populations returned to baseline activity over similar timescales. The function of post-song activity is unclear but may provide a circuit mechanism for birds to rapidly re-engage in song performances given appropriate social feedback or context. Courtship singing is tightly coupled to social interaction with female birds and it is common for male birds to string two or more song phrases together during courtship song^42^ (see **EDFig. 1 & 6**). Post-song activity could also reflect moments when birds are unable to continue singing due to hyperventilation induced by the rapid respiratory patterns associated with song^43^. To explore this idea, we examined whether the duration of song phrases was correlated with the number of active neurons in the post-song period, but found no correlation (*r*^2^ = 0.03). Although the function of post-song activity is unclear, our results indicate that pre-song activity forecasts impending song and suggest a previously unappreciated role for the HVC_RA_ network in planning or preparing to sing.

### Common Preparatory Activity in Premotor Circuits Across Multiple Species

To examine whether preparatory activity is a common circuit mechanism for the production of birdsong, we recorded HVC and RA neural activity in another songbird species, Bengalese finches (*Lonchura striata domestica*). Motor planning and preparation facilitate the accurate execution of fast and precise movements, which are common to the songs of zebra finches and Bengalese finches, however, syllable sequences in Bengalese finches are less stereotyped than those in zebra finches^44^. Using multi-channel neural recordings in HVC, we identified robust preparatory activity several hundreds of milliseconds prior to song onset (**Figure 4a-g**). The pre-song and song– related multiunit spike rates were significantly above baseline (Wilcoxon signed-rank test after Bonferroni correction, n = 29 MU sites. pre-song: *z* = 4.62, *p* = 0.030; song: *z* = 4.70, *p* < 0.001), whereas the post-song spike rates was not (*z* = 2.17, *p* = 0.089). Pre-song activity increased above baseline -1.47 ± 1.05 s prior to song onset, a timescale that closely matched the timing of peak calcium-event rates in peri-song neurons in zebra finches. The offset timing of post-song activity was 0.42 ± 0.23 s. This indicates that preparatory activity in HVC is a common network motif important for song generation and the onset of precise neural sequences.

HVC contains multiple cell types, including interneurons and at least three different classes of projection neurons^45,46^. Multichannel recordings in HVC provide an important read-out of the network activity prior to song onset but alone are insufficient to assess whether this preparatory activity influences descending cortical pathways involved in song motor control. Therefore, we recorded single unit electrophysiological activity from the downstream targets of HVC_RA_ neurons within the cortical premotor nucleus RA. Projection neurons in RA are tonically active at baseline (**Figure 5a**, “Spontaneous”) and exhibit precise bursts of activity during singing (**Figure 5a**, “Song”), a transition well captured by changes in the coefficient of variation of the inter-spike intervals (CV_ISI_, **Figure 5c**). We measured changes in RA neuron activity in the period just prior to song initiation and just after the conclusion of each song bout (between 0.5 and 2.5 s before/after the first/last song syllable). As expected, we did not find substantial differences in spike rates between non-singing and singing states (not shown) but found that the CV_ISI_ for RA neurons changed significantly during song, reaching higher values during pre-song, song, and post-song epochs as compared to spontaneous activity (**Figure 5d,** *p*<0.005, two-sided K-S tests). This suggests that HVC_RA_ sequences associated with preparation to sing propagate to downstream premotor circuits prior to song onset and that HVC continues to influence descending motor pathways following song cessation.

### Peripheral Preparation to Sing

Pre-song activity could reflect motor planning (changes in network activity independent of changes in the motor periphery) and/or motor preparation that functions to coordinate changes in the motor periphery as birds prepare to sing. Song is a respiratory behavior that is primarily produced during expiration and silent intervals in the song correspond to mini-breaths, which are rapid, deep inspirations^31,47^. How birds plan to sing or prepare the respiratory system to sing is poorly understood, but there is evidence that prior to song onset, oxygen consumption decreases and respiratory rate increases^43^. To explore the time course of changes in respiratory patterns in more detail, we used air sac pressure recordings in singing zebra finches (**Figure 6a-c**). During singing, birds significantly accelerated the respiratory rhythm and marginally shifted towards longer periods of expiration during each cycle (**Figure 6b,** respiratory duration pre-song = 0.38 ±0.04 s (SD), song = 0.18 ±0.03 s, post-song = 0.39 ±0.06 s: *F*(2,10) = 46.63, *p* < 0.001; duty cycle (% of time in expiration) pre-song = 58% ±3 (SEM), song = 57% ±2.5, post-song = 61% ±2.4: *F*(2,10) = 3.56, *p* = 0.07). We found that significant changes in respiration also preceded song onset. Respiratory cycle duration significantly accelerated in the last second prior to song onset with relative decreases of expiratory phases (respiratory duration: *F*(3,15) = 7.67, *p* = 0.02, duty cycle: *F*(3,15) = 4.077, *p* = 0.07; **Figure 6d**). Following song termination, birds immediately returned to longer respiratory cycles but during the first second post-song, they spent more time exhaling compared to inhaling, a behavior likely involved in helping to recover from singing-related hyperventilation (respiratory duration: *F*(2,10) = 0.509, *p* < n.s., duty cycle: *F*(2,10) = 6.553, *p* <0 .01; **Figure 6e**). These changes in respiration during the last second before and first second following song support the idea that HVC_RA_ neurons provide descending motor commands that coordinate transitions between non-vocal and vocal states by coordinating respiratory patterns. The lack of changes at earlier time-points prior to singing also indicate that pre-song activity 1-3 s prior to song onset may reflect motor planning or the decision to sing, rather than respiratory preparation. Together these, findings support the idea that HVC_RA_ neurons could function in aspects of motor planning as well as preparation.

## DISCUSSION

Previous studies suggested that the HVC_RA_ network functions exclusively as a time-keeper, encoding motif-level temporal representations of song via propagation of precisely timed neural sequences^1,4,6,28,34,35^. Central to this view is that the network is active only during singing and hence behaves in only two modes, inactive or propagating neural sequences. Our principal result is that neural sequences emerge as part of orchestrated activity across the network of HVC_RA_ neurons and that this activity is correlated with motor preparation prior to song initiation. Peri-song and pan-song neurons forecast the start of singing. Peri-song neurons are inactive during song, whereas pan-song neurons become heterogeneously active prior to time-locked sequential activity during song performances (**Figures 2 & 3**). Preparatory activity in HVC_RA_ neurons precedes the pre-bout activity observed in other classes of HVC neurons in zebra finches and Bengalese finches (**Figure 4**), suggesting that HVC_RA_ neurons may seed network wide changes in activity^39-41^. The rigid stereotypy of singing behavior enables comparisons from different levels of the nervous system and periphery. We find that preparatory activity in HVC_RA_ neurons drives descending motor commands via RA and motor movements that set the stage for producing song (**Figures 5 & 6**). Because song is a self-initiated, volitional behavior, our findings further indicate that the HVC_RA_ network either functions as a sensitive read-out of the decision to sing or as an integral factor in the decision itself. Finally, previous studies have shown that about half of HVC_RA_ neurons are inactive during song^1,27-29^. We find that approximately half of this network is active at peri-song intervals and propose that one function of HVC_RA_ neurons is to plan and prepare for song performance.

Together, these findings support a simple model for song and neural sequence initiation. Preparatory activity in populations of HVC_RA_ neurons drives descending motor commands via RA and its connections to the ventral respiratory group and syringeal motoneurons in the medulla^48-53^. Given the sparsity of HVC_RA_ neuron activity and the convergence of HVC_RA_ input to RA, it is likely that only population activity, like that described here, is sufficient to drive bursting in RA. These motor signals increase respiratory rate, bringing it closer to the high rate needed to coordinate production of song syllables. Because the initiation of singing requires precise coordination between respiratory state and descending motor commands, we hypothesize that recurrent projections from the brainstem update activity in HVC and trigger the initiation of neural sequences once the periphery is readied for the first respiratory cycle for song^29,54,55^. Circuitry related to recurrent projections into HVC increase the HVC_RA_ activity hundreds of milliseconds prior to song onset^56-58^ and thus may also mediate shifts in activity between peri-song and pan-song neurons.

Absent from this model is how activity in peri-song and pan-song neurons is first initiated seconds before song onset. HVC receives input from cholinergic neurons in the basal forebrain^59-61^, noradrenergic neurons in the locus coeruleus^62^, and dopaminergic neurons in the midbrain^63^, any or all of which could play potent roles in shifting the excitability of subsets of HVC_RA_ involved in song preparation and initiation. Also absent are specific predictions about the role of peri-song neurons and pan-song neurons in motor-planning. It is not yet known whether preparatory activity in HVC is necessary for song initiation or production. These experiments will require novel closed-loop manipulations to exclusively disrupt peri-song or pan-song neuron activity independent of any activity involved in neural sequences. If possible, such experiments will undoubtedly lend insights into whether HVC is involved in aspects of motor-planning independent of motor-preparation.

Sequential activation of neurons is thought to provide computational advantages for encoding temporal information associated with episodic memories or behavioral sequences. Neural sequences in HVC provide one of the cleanest examples for linking brain activity with a naturally learned and volitionally produced skilled motor behavior. Our study provides a glimpse of how these sequences emerge through temporally coordinated transitions within a potentially hierarchically organized network and suggests a general framework for initiating the production of skilled motor behaviors.

## Supporting information

## Acknowledgments

The authors thank Joseph Takahashi for generous use of a miniscope for calcium imaging experiments, J Holdway, M Ikeda, D Merullo, B Pfeiffer, A Wood, and L Xiao for comments on the manuscript and discussions, the Genie Project at Janelia for development of calcium indicators, T Komiyama and A Peters for analysis codes, J Holdway and A Guerrero for laboratory support and animal husbandry, and J Dukes for assistance with data analysis. This research was supported by grants from the US National Institutes of Health R01NS108424 and R01DC014364 to TFR, R01NS108424 to TFR, RHRH and BGC, and R01NS084844 and R01NS099375 to SJS, the National Science Foundation IOS-1457206 to TFR, and the Swiss National Science Foundation 31003A_127024 and 31003A_156976 or RHRH.

## Author contributions

V.K.D. and T.F.R. initiated this study. V.K.D. collected and analyzed calcium imaging data. B.G.C. collected and analyzed respiratory data. R.O.T., S.K. and R.H.R.H. collected and analyzed the chronic recording data from HVC. S.J.S. collected and analyzed the chronic recording data from RA. V.K.D. and T.F.R. wrote the paper with input from all authors.

## Author Information

The authors declare no competing financial interests. Correspondence and requests for materials should be addressed to T.F.R.

## Methods

### Animals

Experiments described in this study were conducted using adult male zebra finches and Bengalese finches (>90 days post hatch). During experiments, birds were housed individually in sound-attenuating chambers on a 12/12 h day/night schedule and were given ad libitum access to food and water. All procedures were performed in accordance with established protocols approved by Animal Care and Use Committee’s at UT Southwestern Medical Centers, Texas Christian University, Emory University, and the Korea Brain Research Institute.

### Viral Vectors

The following adeno-associated viral vectors were used in these experiments: AAV2/9.CAG.Flex.GCaMP6s.WPRE.SV40 (University of Pennsylvania) and AAV2/9.CMV.Cre.WPRE.SV40 (University of Pennsylvania). All viral vectors were aliquoted and stored at -80°C until use.

### Imaging

Two-photon microscopy was conducted with a commercial microscope (Ultima IV, Bruker) running Prairie View software using a 20× (1.0 NA) objective (Zeiss) with excitation at 920 nm (Ti-Sa laser, Newport). Imaging was conducted in lightly anesthetized animals that were head-fixed using a custom-built apparatus. Single-photon images of HVC were acquired through the cranial window using a sCMOS camera (QImaging, optiMOS). These images were used to guide placement of the baseplate for miniaturized single-photon microscope.

### Stereotaxic Surgery, Cranial Windowing, and Baseplate Implantation

All surgical procedures were performed under aseptic conditions. Birds were anesthetized using isoflurane inhalation (^~^1.5-2%) and placed in a stereotaxic apparatus. Viral injections were performed using previously described procedures (Roberts et al., 2012, 2017) at the following approximate stereotaxic coordinates relative to interaural zero and the brain surface (rostral, lateral, depth, in mm): HVC (0, 2.4, 0.1-0.6); and RA (−1.0, 2.4, 1.7-2.4). The centers of HVC and RA were identified with electrophysiology. For calcium imaging experiments, 1.0 – 1.5 μL of Cre-dependent GCaMP6s was injected at 3 different sites into HVC and 350 nL of Cre was injected into RA. Viruses were allowed to express for a minimum of 6 weeks before a cranial window over HVC was made. Briefly, a unilateral square craniotomy (^~^3.5 × 3.5 mm) was created over HVC and the dura was removed. A glass coverslip was cut to match the dimensions of the craniotomy and held in place with a stereotaxic arm as Kwik Sil was applied to the edges of the cranial window. Dental acrylic was applied over the Kwik Sil and allowed to slightly overlap with the glass coverslip to ensure the window would not move and would apply the appropriate amount of pressure to the brain. An aluminum head post was affixed to the front of the bird’s head to enable head-fixed imaging under the 2-photon microscope and to enable head-fixation for baseplate implantation. Following verification of labeling, identification of HVC boundaries, and high-resolution images of neurons under the 2-photon microscope, the bird was lightly anesthetized with isoflurane and the miniaturized fluorescent microscope (Inscopix) was placed over the cranial window. The field of view that matched the 2-photon images was identified and the focal plane that enabled the largest number of neurons to be in focus was selected. Dental acrylic was used to fix the baseplate in the desired position and any exposed skull was covered with dental acrylic. Once the dental acrylic dried, the microscope was removed from the baseplate and the bird was allowed to recover overnight. About 30 minutes before the birds’ subjective daytime, the microscope was attached to the counterbalance (Instech) with enough cable to allow the bird to move freely throughout the cage. The microscope was then secured to the baseplate with a setscrew. The bird was allowed to wake up and accommodate to the weight of the microscope over the next 2-3 days.

### GCaMP6s Imaging Using a Miniaturized Fluorescent Microscope

The miniaturized fluorescent microscope (Inscopix) was not removed following successful baseplate implantation and remained attached to the birds’ head until either the cranial window closed or 7-10 days had passed. The counterbalance was adjusted based on the observed behavior of the bird and its ability to move freely. The female was not housed in the cage with the male bird, but instead was introduced to the males during a minimum of 3 morning and afternoon sessions to evoke directed song. Video recording was first started followed by 5 to 10 seconds of spontaneous recording with the miniaturized fluorescent microscope. The female bird was placed in the cage as quickly and as with little disruption as possible for each session. If the male bird did not sing within a minute of the females’ presence, the session was stopped, and the female was removed. All trials were recorded on video, and audio was recorded using Sound Analysis Pro (SAP) software and the HD video camera microphone. Calcium imaging was performed at 30 frames per second (fps), at 1080×1920 resolution, Gain was set to 4, and Power was set to 90% for all birds, behavioral videos were collected at 24 fps. Calcium imaging data and behavioral data was synchronized using start of calcium imaging on a frame by frame basis.

### Defining Song and Peri-Song Behavior

We defined singing as the time from the first introductory note to the end of the last syllable of the birds’ bout. The time (inter-bout interval) between multiple renditions of song motifs determined whether subsequent singing was included as another bout in the phrase or the start of another phrase. Silent gaps greater than 1 second but less than 2 seconds between the offset of the last syllable and the start of the next syllable in a motif were treated as inter-bout intervals and calcium events occurring during this time were excluded from song-activity analysis. Inter-bout intervals less than 2 seconds were included in the same phrase. Inter-bout intervals greater than 2 seconds served as the cutoff between two separate phrases. Calcium events occurring during inter-phrase intervals (time between end of previous song syllable and start of next song syllable, must be greater than 2 seconds) were correlated with the closest phrase onset or offset.

### Calcium image processing and analysis

Calcium images were collected using the miniaturized fluorescent microscope developed by Inscopix^32^. First, the FOV was spatially cropped to exclude pixels that did not include neurons or observable changes in fluorescence. Next, the preprocessing utility within the Mosaic data analysis software was used to spatially bin the images by a factor of 2 to reduce demands on computer memory and enable faster data processing. The TurboReg implementation within Mosaic was used to perform motion correction. A reference image was created using a maximum intensity projection of the dataset and the images were aligned in the x and y dimension to the reference image. Imaging datasets with translational motion greater than 20 pixels in either the x or y dimensions were excluded from further data analysis. Post-registration black borders were spatially cropped out. The resulting spatially-cropped, preprocessed, and motion corrected calcium imaging datasets were exported for further analysis in custom Matlab scripts.

We performed ROI-based analysis on the motion-corrected calcium imaging datasets using previously described methods^3^. ROIs were manually drawn around identifiable soma and a secondary ROI that extended 6 pixels around the boundaries of the neuronal ROI was used to estimate background fluorescence (i.e. neuropil or other neurons). The pixel values were averaged within the neuronal and background ROIs, and background fluorescence signal was subtracted from neuronal signal. An iterative procedure using custom Matlab scripts were used to estimate baseline fluorescence, noise, and active portions of the traces^3^. A subset of calcium images were re-analyzed using previously described constrained non-negative matrix factorization (CNMF) methods, but calcium fluorescence traces were identical to the traces pulled out by the ROI-based analysis^64^. Calcium traces generated by ROI-based analysis were further deconvolved to produce inferred calcium traces using the pool adjacent violators algorithm (PAVA)^65^. The deconvolved calcium traces were normalized to values between 1 and 0 to enable visualization of activity across different neurons during the same trial. Calcium transients that were 3 SD. above baseline activity were recorded as events. The corresponding onset times and the rise times to peak fluorescence of individual calcium transients were correlated with synchronized behavior.

All calcium events were first categorized as falling into peri-song or song behavioral epochs. Peri-song was limited to the 5 second period before song onset (including introductory notes) and the 5 second period after offset of the last syllable. These event counts were used to assign a Phrase index to all imaged neurons. Neurons that had fewer than 2 calcium events recorded over a day of singing were excluded from further analysis because sparsity of calcium events could spuriously identify neurons as peri-song or song exclusive. We combined the number of calcium events from neurons imaged across multiple trials during the same day. Neurons imaged across multiple days were treated as unique neurons. The phrase index was calculated as a ratio of the total number of song events imaged from a neuron during a day subtracted by the number of peri-song events to the total number of calcium events. This bounded the phrase index to values of -1 (peri-song exclusive) and +1 (Song exclusive). We used the phrase index to examine the timing properties of neurons active only during peri-song, song, or both behavioral epochs.

To examine the distribution of calcium events, we generated histograms with bin sizes of 200 ms. Calcium event onset times were shifted to negative values if they occurred prior to song onset. Calcium events occurring after song offset were shifted by 1 second to allow visualization of a 1 second song phrase window. The number of active neurons was calculated in 200ms bins, if a neuron had more than 1 calcium event within a 200 ms window it was considered active only once. Event rates were calculated as a moving average of the number of events organized in 100ms windows before and after song onset. Rates were calculated by phrase using a 1 second moving average window, this window reaches a minimum of 500 ms at boundaries (−5, +5 seconds, and at song onset and offset). The average event rate and standard deviation was calculated from all pre and post phrase event rates from all birds.

### Fluorescence Analysis Across Intervals

Average fluorescent changes were measured for each neuron across baseline, pre-song, post-song, and song behavioral periods. Baseline was defined as a behaviorally quiet period covering 5s of fluorescent activity that was ≥10 s removed from periods of singing or calling. Pre-song was 5s before phrase onset and post-song was 5s after phrase offset. The background subtracted fluorescent traces were used to measure average fluorescence across the above intervals for all phrases and all birds. Averaged fluorescent values were than normalized to the average fluorescence measured during song.

### Comparison of SNR and Calcium Event Peak Magnitudes

We measured the SNR of events occurring during song for a subset of pan-song and song neurons. The SNR was calculated as a ratio of peak fluorescence for each song event per neuron to the average fluorescence from baseline period within the trial (as above, 5s of fluorescent activity that was ≥10s removed from periods of singing or calling). We determined the average SNR for each neuron and examined differences between pan-song and song neurons.

Peak magnitudes during peri-song and song periods of pan-song neurons were measured using normalized deconvolved fluorescent traces. The peak values for each pan-song neuron (with phrase indices between -0.18 to +0.18) during peri-song and song periods were used to evaluate potential differences between calcium events occurring outside of song versus during song.

### Neural Recordings

Multiunit recordings of HVC neurons were collected from three adult (>90 days old) male Bengalese finches. All procedures were approved by the Korea Brain Research Institute. An array of 16 tungsten microwires (175 μm spacing, OMN1005-16, Tucker Davis Technologies) was implanted into left HVC. The location of HVC was identified by searching for spontaneous spike bursts and for antidromic response to stimulation in RA. The extracellular voltage traces of all channels from birds singing alone (without presentation of female) were amplified and recorded with an interface board (RHD2132, Intan Technologies) at a sampling rate of 25kHz. The interface board was tethered to a passive commutator (Dragonfly Inc.) via a custom-made light-weight cable. In total we obtained HVC recordings from 35 electrode channels in three birds (15, 7, and 8 channels, respectively), all of which showed spontaneous bursts typical of HVC neurons. The impedance of successful electrodes was around 100-300 kΩ.

Recorded signals were bandpass filtered (0.3-5kHz) and negative signal peaks exceeding 4 SD of spontaneous activity (no-song period separated more than 10 seconds from nearest song bouts) were interpreted as multi-unit spikes. In total 38, 62, and 212 song onsets, and 42, 23, and 280 offsets were identified in these birds, respectively. We produced firing rate-traces from each electrode channel with 10ms resolution and averaged them across song renditions. After smoothing by moving average with 500ms window, the averaged firing rates were normalized into 0 to 1 as spanning between respective mean firing rates during spontaneous activity and singing (0-3s after song onset) to remove any bias among channels to obtain the general trend of onset-and offset-related firing across channels and birds.

The significance of activity elevation during pre-song, song, and post-song periods from the spontaneous activity level was tested by Wilcoxon signed-rank test with significant level at 0.05 after Bonferroni correction for multiple comparison.

Onset of pre-song and offset of post-song activity were estimated for each channel as the smoothed spike rate trajectory was exceeded a threshold which was defined as mean + 2 SD of the spontaneous spike rate.

Single-unit and multiunit recordings of RA neurons were collected from six adult (>140 days old) male Bengalese finches as described previously^66^. All procedures were approved by the Emory University Institutional Animal Care and Use Committee. Briefly, an array of four or five high-impedance microelectrodes was implanted above RA. We advanced the electrodes through RA using a miniaturized microdrive to record extracellular voltage traces as birds produced undirected song (i.e., no female bird was present). We used a previously described spike sorting algorithm to classify individual recordings as single-unit or multiunit^67^. In total, we recorded 19 single units (multiunit recordings were not analyzed further in this study). Based on the spike waveforms and response properties of the recordings, all RA recordings were classified as putative projection neurons^67-69^.

### Analysis of Chronic Recording Data

To analyze the variation in inter-spike-interval (ISI) in different time periods (Figure 5), we restricted our analysis to cases in which we collected at least one recording that included the relevant song epoch. “Spontaneous” epochs were sampled from neural activity recorded more than 10 sec after the nearest song bout. “Pre-song” activity was sampled from between 2.5 and 0.5 s prior to the first song syllable or introductory note. “Song” activity was sampled from the onset of the first song syllable until the offset of the last syllable in a bout. “Post-song” activity was sampled from between 0.5 and 2.5 s after the offset of the last syllable in a bout. In some cases, we did not have sufficient data available from all epochs for all neurons (note the variation in the number of neurons included in the analyses shown in Fig. 5d).

### Air Sac Recording Procedures

Subsyringeal air pressure was recorded from six birds in directed singing conditions. Directed song was defined as a female presented in an adjacent cage during a two-hour recording period. Data from four of the birds were re-analyzed from a previously published study (Cooper & Goller, 2006) and data from two additional birds were collected to replicate the effects observed in the previously collected data70. As described in (Secora et al. 2004), each bird was accustomed to carrying a pressure transducer that was held in place on the bird’s back with an elastic band^71^. To facilitate relatively free lateral and vertical movement in the cage, the weight of the transducer was offset by a counter-balance arm. Subsyringeal air pressure surgery was performed after birds sang while carrying the pressure transducer. Prior to insertion of the air pressure cannula, animals were deeply anesthetized as verified by an absence of a toe-pinch response. A small opening in the body wall below the last rib was made with a fine pair of micro-dissecting forceps, and a flexible cannula (silastic tubing, OD 1.65 mm, 6.5 cm length) was inserted into the body wall and suture was tied around the cannula and routed between the 2^nd^ and 3^rd^ ribs to hold it in place. The skin was sealed to the cannula with tissue adhesive. The free end of the cannula was attached to the pressure transducer. This allowed for measurement of relative subsyringeal air pressure changes inside the thoracic air sac before, during, and after spontaneously generated song events. Birds were monitored following surgery until they perched in the recording chamber.

The voltage output of the pressure transducer was amplified (50-100 x) and low-pass filtered (3 kHz cutoff; Brownlee, Model 440, Neurophase, Santa Clara, CA). Respiration was recorded for five seconds prior to and following singing epochs using a National Instruments analog-to-digital conversion board (NI USB 6251, Austin, TX) controlled by Avisoft Recorder software (Avisoft Bioacoustics, Berlin, Germany). Data were collected in wav file format, 16 bit resolution, with sampling rates varying from 22.05 to 40 kHz. Songs were selected for analysis that contained at least 3 s of uninterrupted quiet respiration prior to and following song. Songs that were preceded by calls, drinking, defecation, or movement-related activity were excluded from the analysis.

### Air sac data classification

Air pressure was analyzed as respiratory cycles, which was defined as an inspiration followed by expiration. The onset of inspiration was identified as subambient air pressurization and the return to ambient pressure following the expiratory phase of the cycle. The cycle duration (s), duty cycle (% time spent in the expiratory phase of respiration), and average rectified amplitude (a.u.) was calculated for each cycle. Song respiration was analyzed prior to song onset and following song termination. Song onset was defined as the inspiration preceding the first introductory note; using this marker, the onset time for each respiratory cycle in the pre-song recording period was determined. The conclusion of song was defined as the termination of the expiration generating the last song syllable in the bird’s song bout. The timing of the respiratory cycles following song were identified relative to the song termination marker.

### Statistical analyses of respiratory data

For statistical analyses of the respiratory data, each bird contributed a single average value for each measured parameter (cycle duration, duty cycle, average amplitude). A repeated measures ANOVA was used to determine how respiration changes prior to and following song. For each bird, ten to twenty songs were identified for the statistical analysis (see above for criteria). The average for the pre-and post-song (3-5s) for each measured parameter for each bird was calculated. To evaluate the time course of change in respiratory patterns preceding and following song, the average for one second bins for each bird were used in the repeated measures ANOVA. In cases where the assumption of sphericity was violated, the Greenhouse-Geisser correction for the *degrees of freedom* was used. All *p* values reported are based on this correction. An *a priori* alpha level of .05 was used for determining statistical significance.

