## Supplementary material for "Transitioning between preparatory and precisely sequenced neuronal activity in production of a skilled behavior"

### Extended Data Table 1

| Bird | Trial | Type | Syllables | Motifs | Bouts | Phrases | Singing Duration (s) |
|---|---|---|---|---|---|---|---|
| O262 | 132226 | Directed | 30 | 6 | 3 | 1 | 4.92 |
| | 135632 | Directed | 25 | 5 | 1 | 1 | 4.16 |
| | 142315 | Directed | 15 | 3 | 1 | 1 | 2.43 |
| | 173107 | Directed | 25 | 5 | 1 | 1 | 3.29 |
| | 174745 | Directed | 25 | 5 | 2 | 1 | 4.08 |
| | 165515 | Directed | 25 | 5 | 1 | 1 | 4.54 |
| | 170812 | Non-Singing* | N/A | N/A | N/A | N/A | N/A |
| O248 | 162048 | Directed | 244 | 49 | 24 | 10 | 31.73 |
| | 162201 | Directed | 85 | 17 | 7 | 5 | 10.58 |
| | 160046 | Directed | 170 | 34 | 21 | 4 | 22.92 |
| | 152940 | Directed | 25 | 5 | 1 | 1 | 3.625 |
| | 140745 | Undirected | 60 | 12 | 3 | 2 | 8.68 |
| | 141006 | Undirected | 100 | 20 | 4 | 2 | 15.952 |
| | 141059 | Undirected | 130 | 26 | 4 | 2 | 19.918 |
| | 131612 | Undirected | 110 | 22 | 4 | 3 | 16.83 |
| | 132211 | Non-Singing | N/A | N/A | N/A | N/A | N/A |
| | 131904 | Non-Singing | N/A | N/A | N/A | N/A | N/A |
| | 123106 | Non-Singing** | N/A | N/A | N/A | N/A | N/A |
| O213 | 164915 | Directed | 30 | 7 | 4 | 1 | 5.1 |
| | 170209 | Directed | 27 | 6 | 3 | 1 | 4.499 |
| | 163439 | Directed | 27 | 6 | 3 | 1 | 4.55 |
| | 165959 | Directed | 12 | 3 | 2 | 1 | 2.07 |
| Y204 | 151708 | Directed | 30 | 9 | 2 | 1 | 5.61 |
| | 155748 | Directed | 14 | 4 | 1 | 1 | 2.57 |
| B89 | 162205 | Directed | 26 | 8 | 1 | 1 | 4.68 |
| B263 | 163144 | Directed | 21 | 3 | 2 | 1 | 7.34 |
| | 162125 | Directed | 14 | 2 | 1 | 1 | 2.72 |
| | 140432 | Directed | 28 | 4 | 2 | 1 | 4.4 |
| | 135424 | Non-Singing*** | N/A | N/A | N/A | N/A | N/A |
| **TOTALS** | **29** | | **1298** | **266** | **98** | **45** | **197.194** |

**Table 1**. Summary of behavioral data set for *in vivo* calcium imaging experiments. *Male did not sing despite having a female present. **Male was actively calling during this trial. ***Male did not sing despite being in the presence of a female, however, the bird does perform introductory notes.

### Extended Data Figure 1

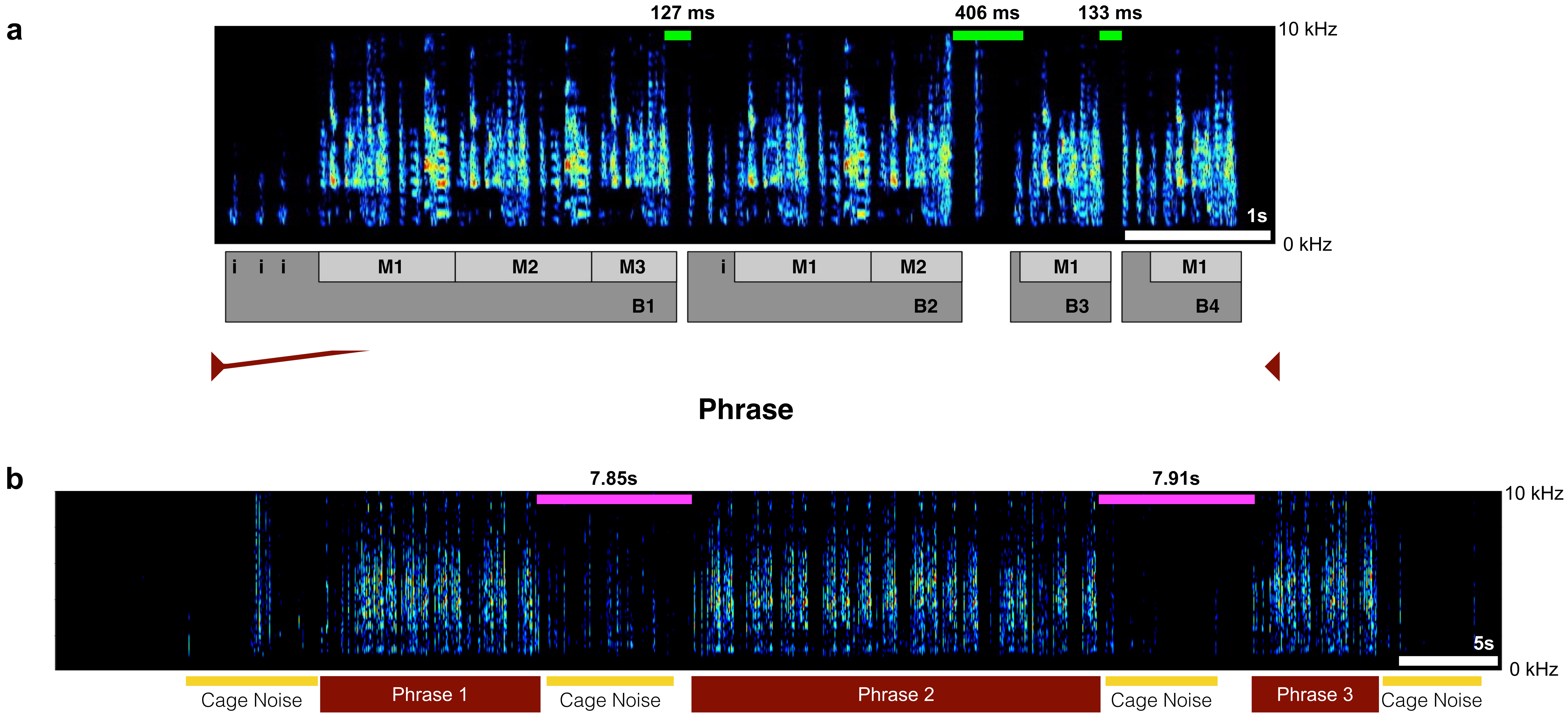


**Figure 1**. a) A spectrogram of a single song phrase composed of 4 bouts and 7 motifs (Bout 1: 3 motifs, Bout 2: 2 motifs, Bout 3: 1 motif, Bout 4: 1 motif). Green bars on top of spectrogram indicate representative silent periods between bouts and numbers above indicate duration in milliseconds. b) A spectrogram highlighting a trial where the bird sang 3 different phrases. The magenta bars above the spectrogram indicate silent periods between phrases and the numbers above indicate duration in seconds. Yellow bars below spectrogram indicate cage noise.

### Extended Data Figure 2

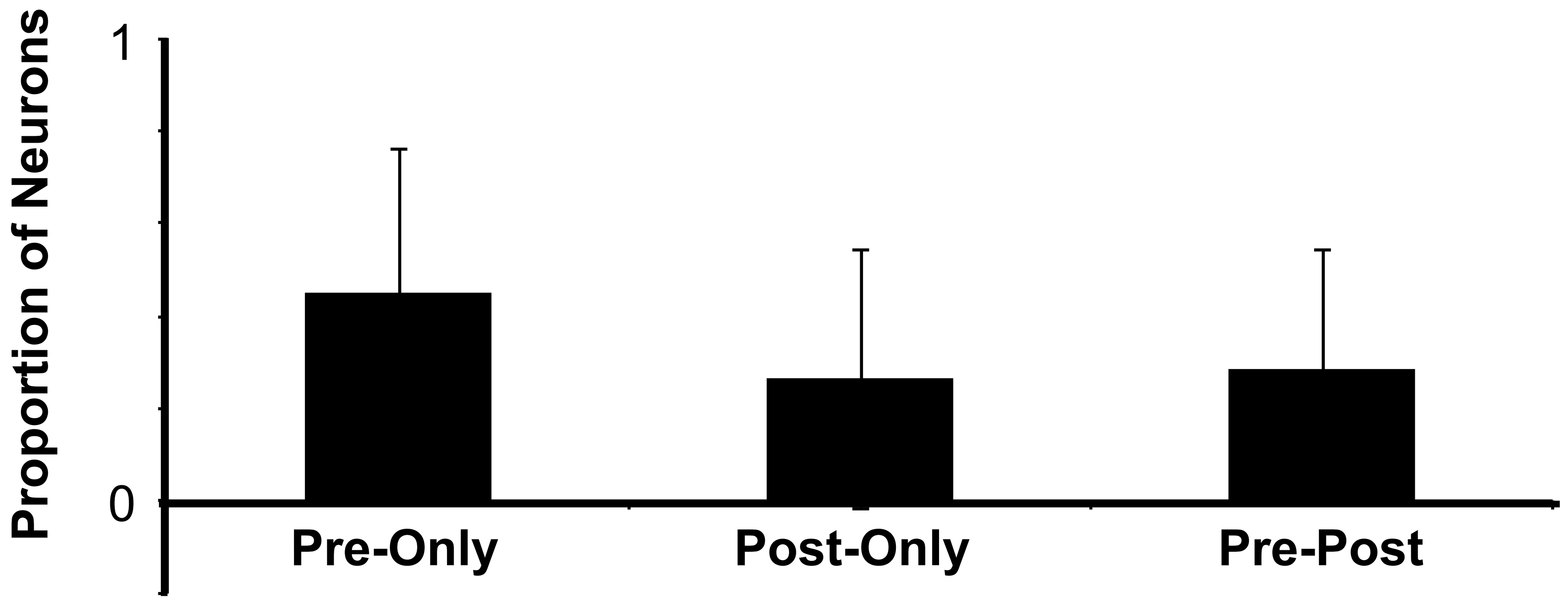


**Figure 2**. Proportion of imaged neurons by bird (N = 6 birds, 197 neurons) that exhibited calcium events only before song onset calcium events (Pre-only, 0.44 ± 0.31, SD), only after song offset calcium events (Post-only, 0.26 ± 0.28), or were active before and after song (Pre-Post, 0.29 ± 0.25).

**Extended Data Figure 3**

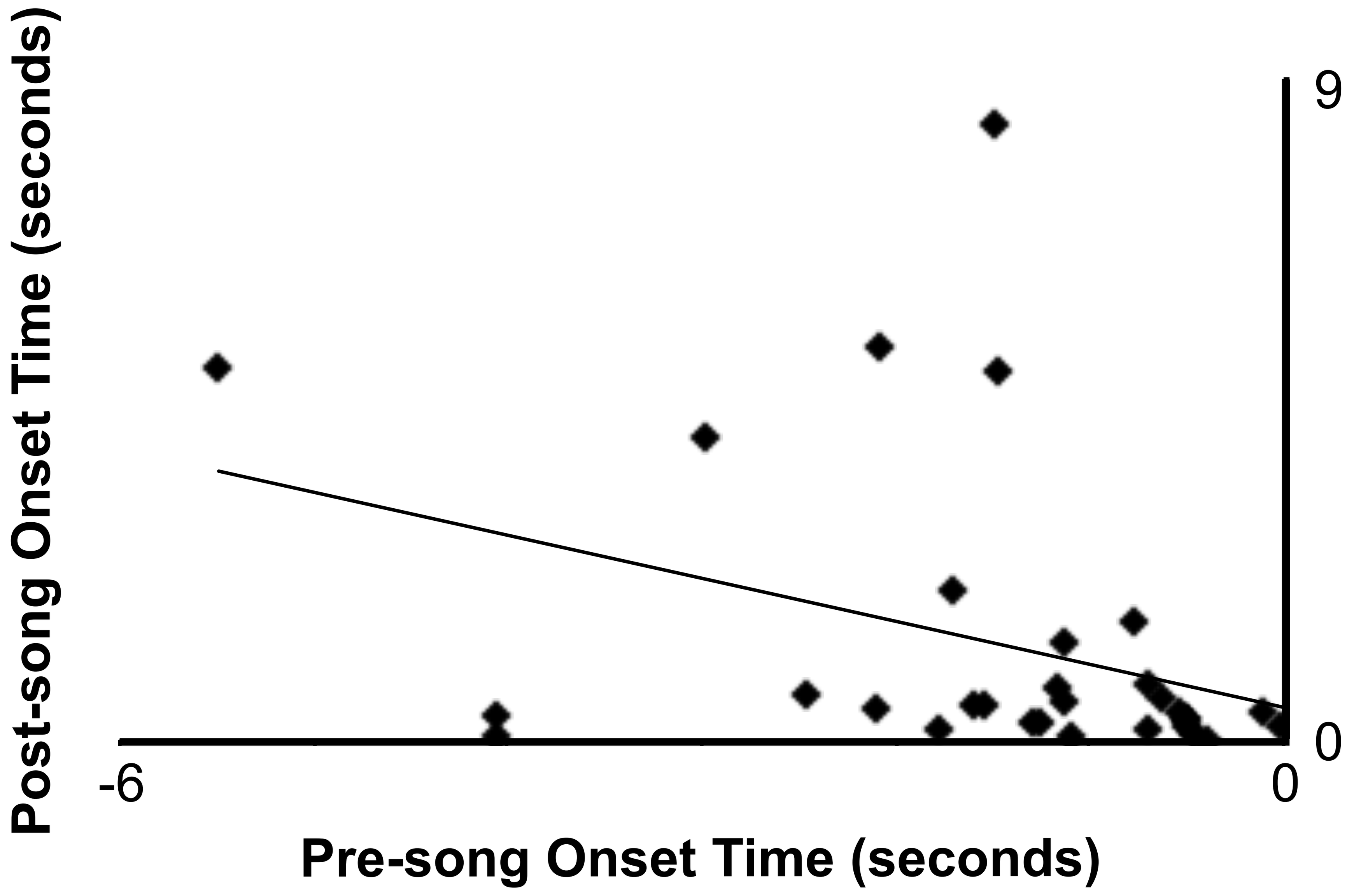


**Figure 3**. 62 events (31 paired pre-song and post-song events) from 31 neurons from 6 birds. Showing a small positive correlation ($r^2$ = 0.127) between pre-song onset time to post-song onset time in cases where a neuron was active only once before and after singing.

**Extended Data Figure 4**

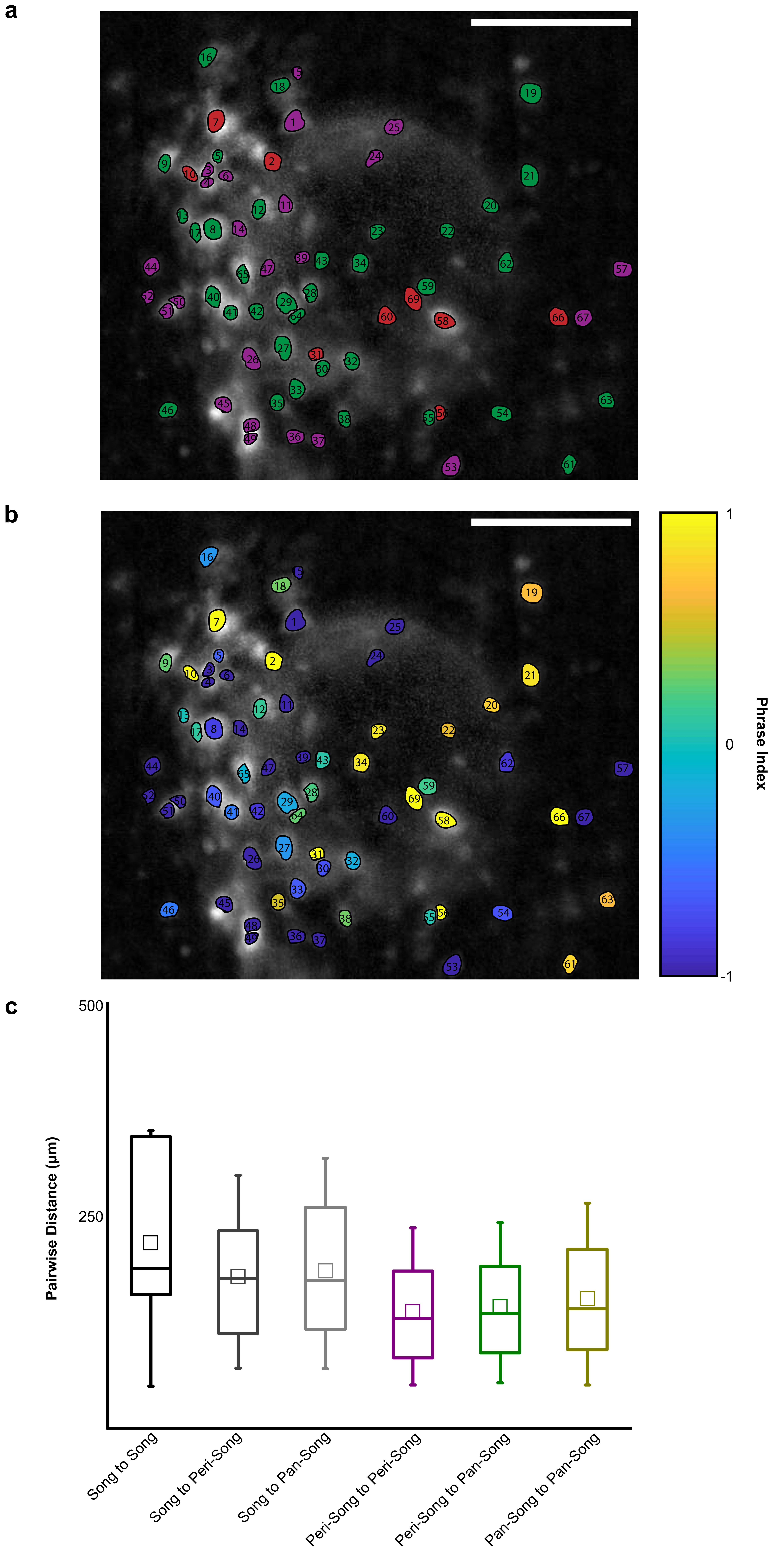


**Figure 4**. a) FOV of ROIs identified in one exemplary bird (O262). Peri-song neurons are shown in purple, pan-song neurons are shown in green, and song neurons are shown in red. Scale bar is 100 micrometers. b) Same FOV as in a, but neurons are color coded by their phrase index. c) Euclidean distances between neurons shown in figure a. Distances (in micrometers) were calculated between the three functional neuron pools mentioned in this paper: peri-song neurons, pan-song neurons, and song neurons. Song-song to peri-peri, p=0.02, song-song to peri-pan, p = 0.04, Mann-Whitney U Test. 2278 pairwise distances.

### Extended Data Figure 5

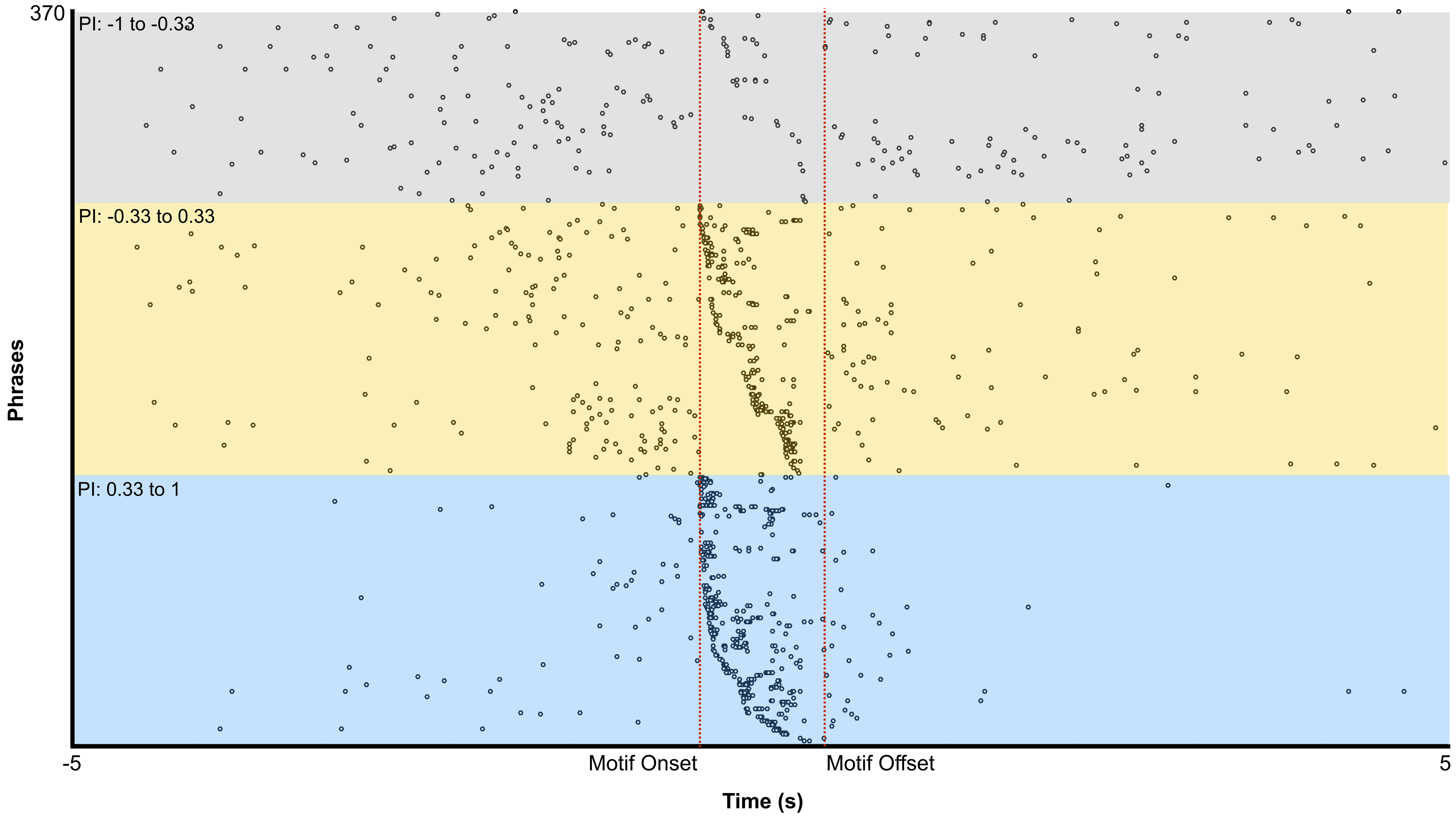


**Figure 5**. Pan-song neuron events organized in tripartite groups according to phrase index. Within each subdivision neurons are organized by Motif onset times. 23 neurons are shown between -1 to -0.33. 56 neurons are shown between -0.33 to 0.33. 53 neurons are shown between 0.33 to 1.

### Extended Data Figure 6

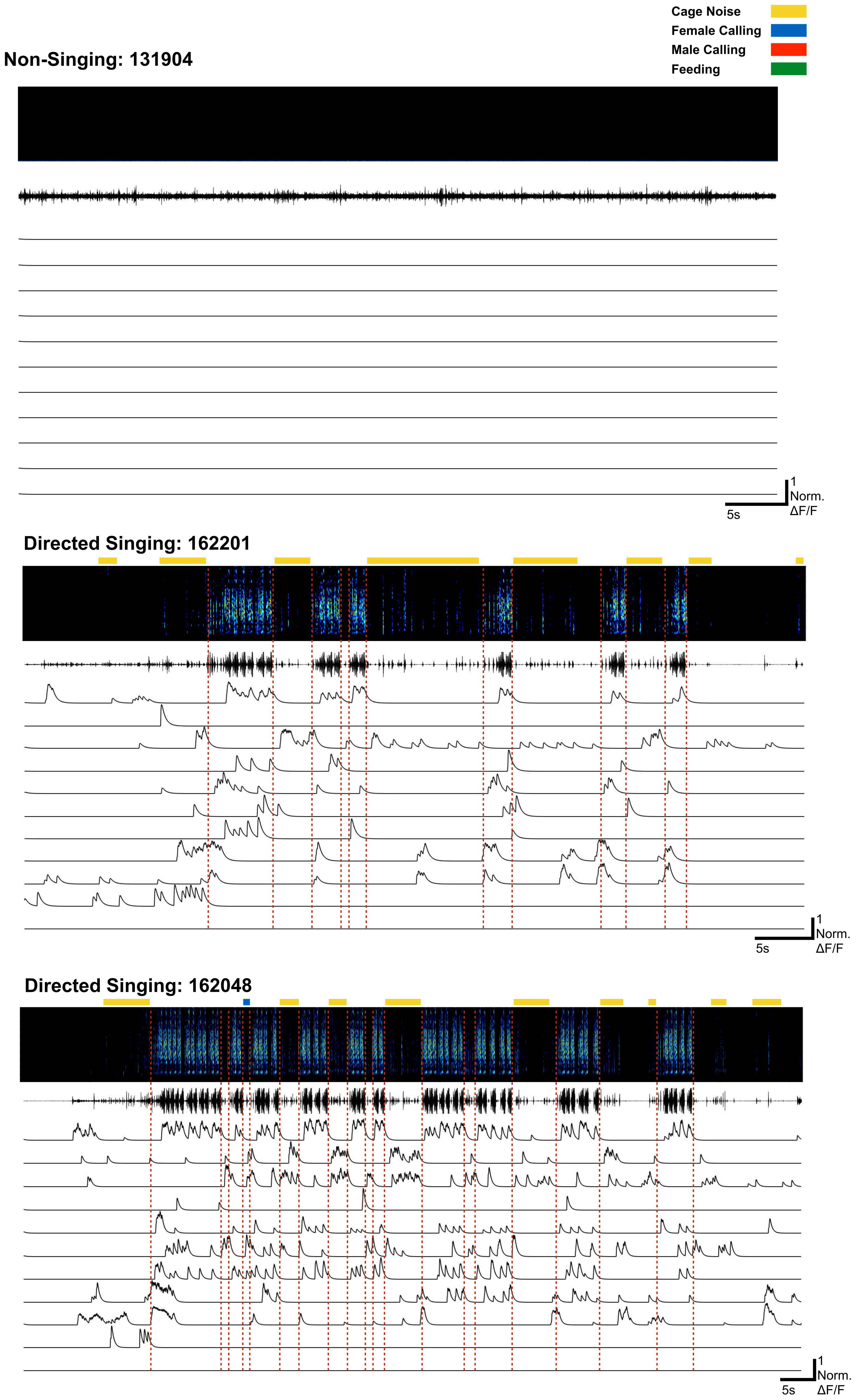

**Extended Data Figure 6**

**Directed Singing: 160046**

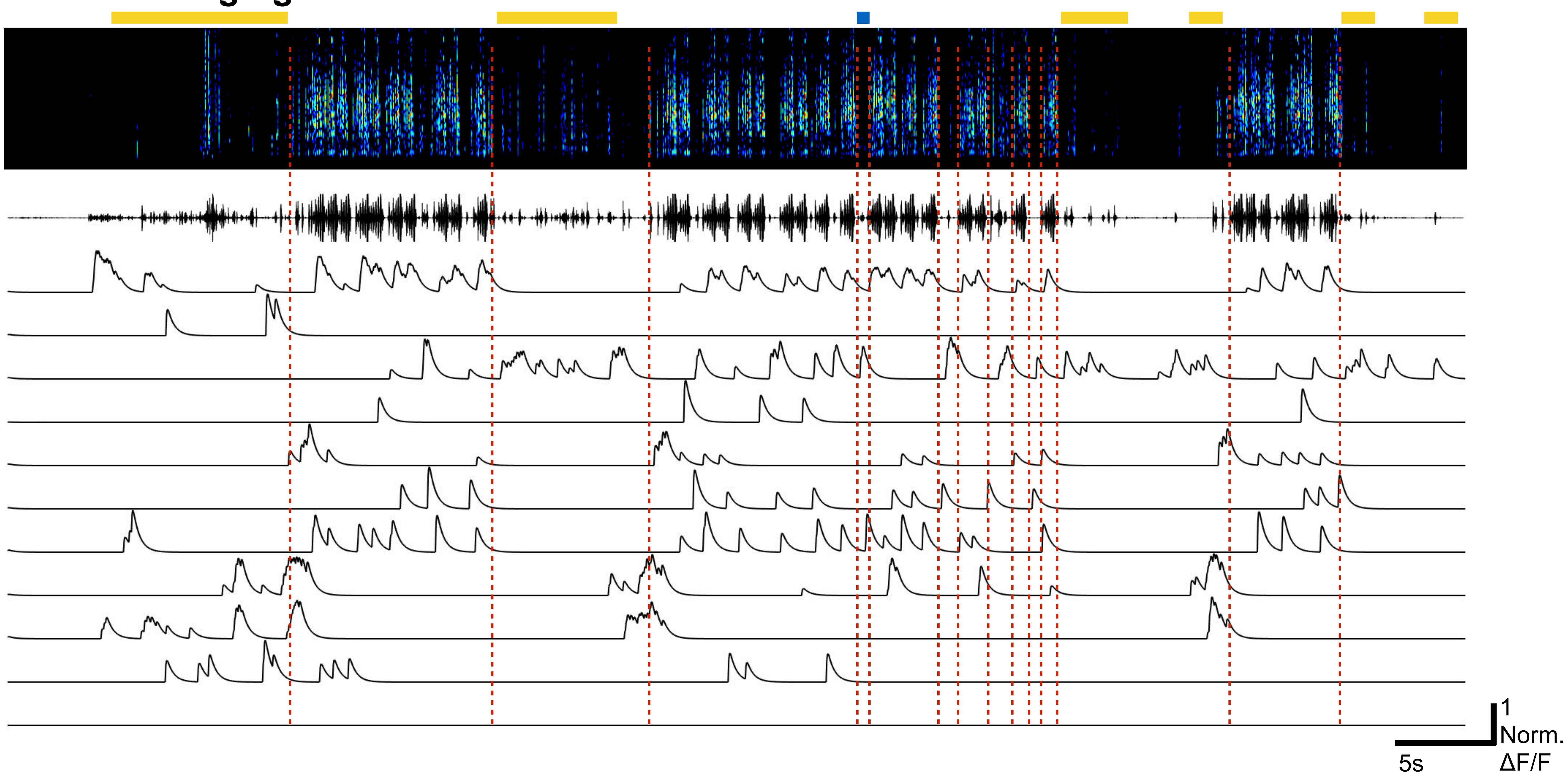


**Directed Singing: 152940**

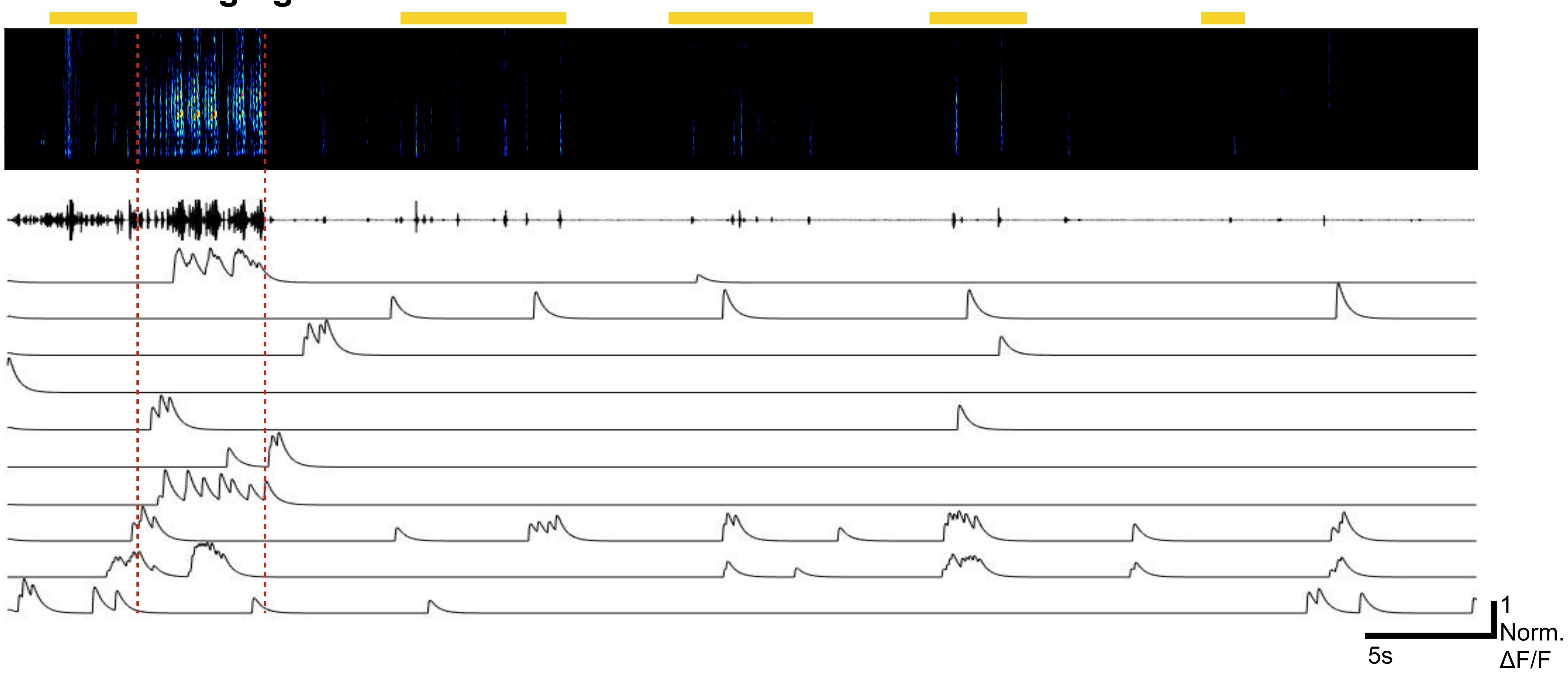


**Undirected Singing:141059**

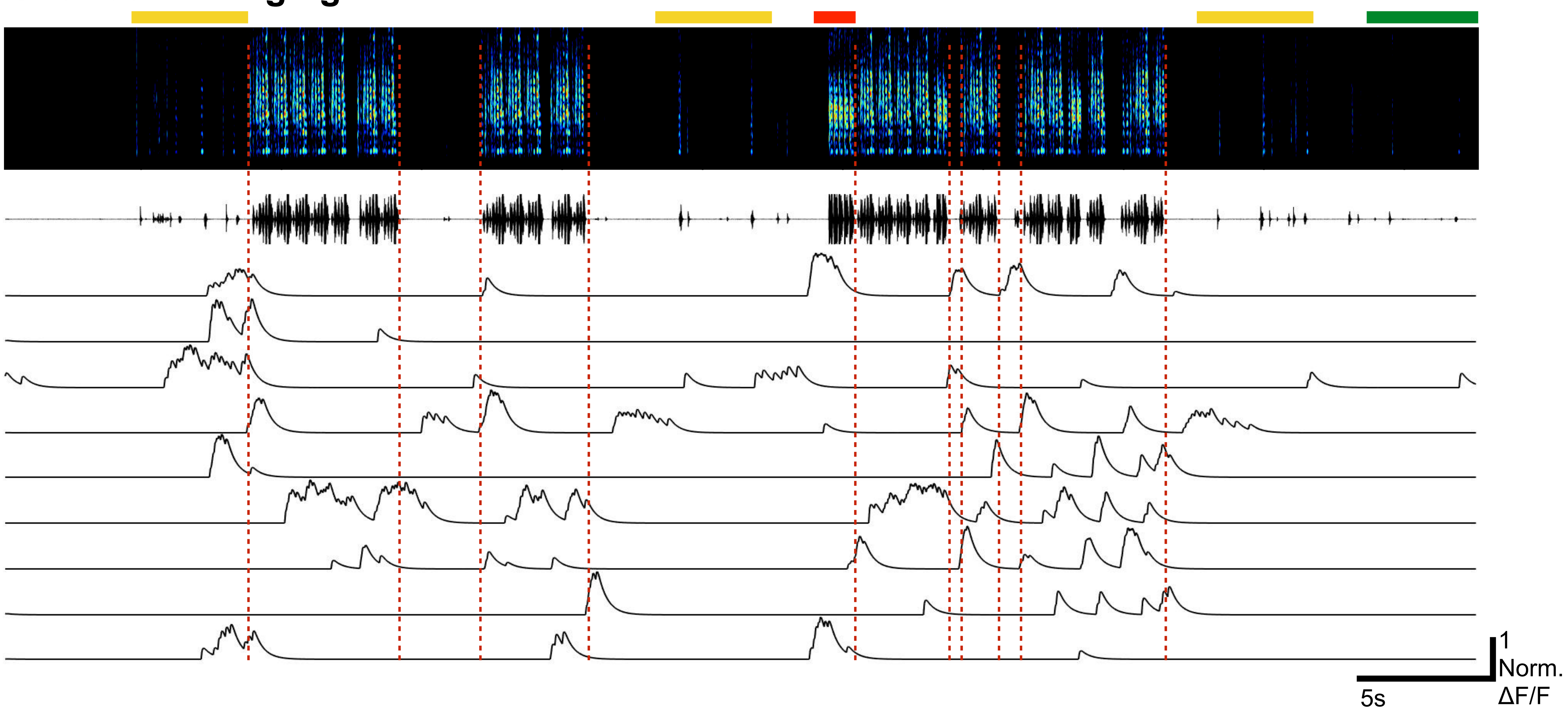

**Extended Data Figure 6**

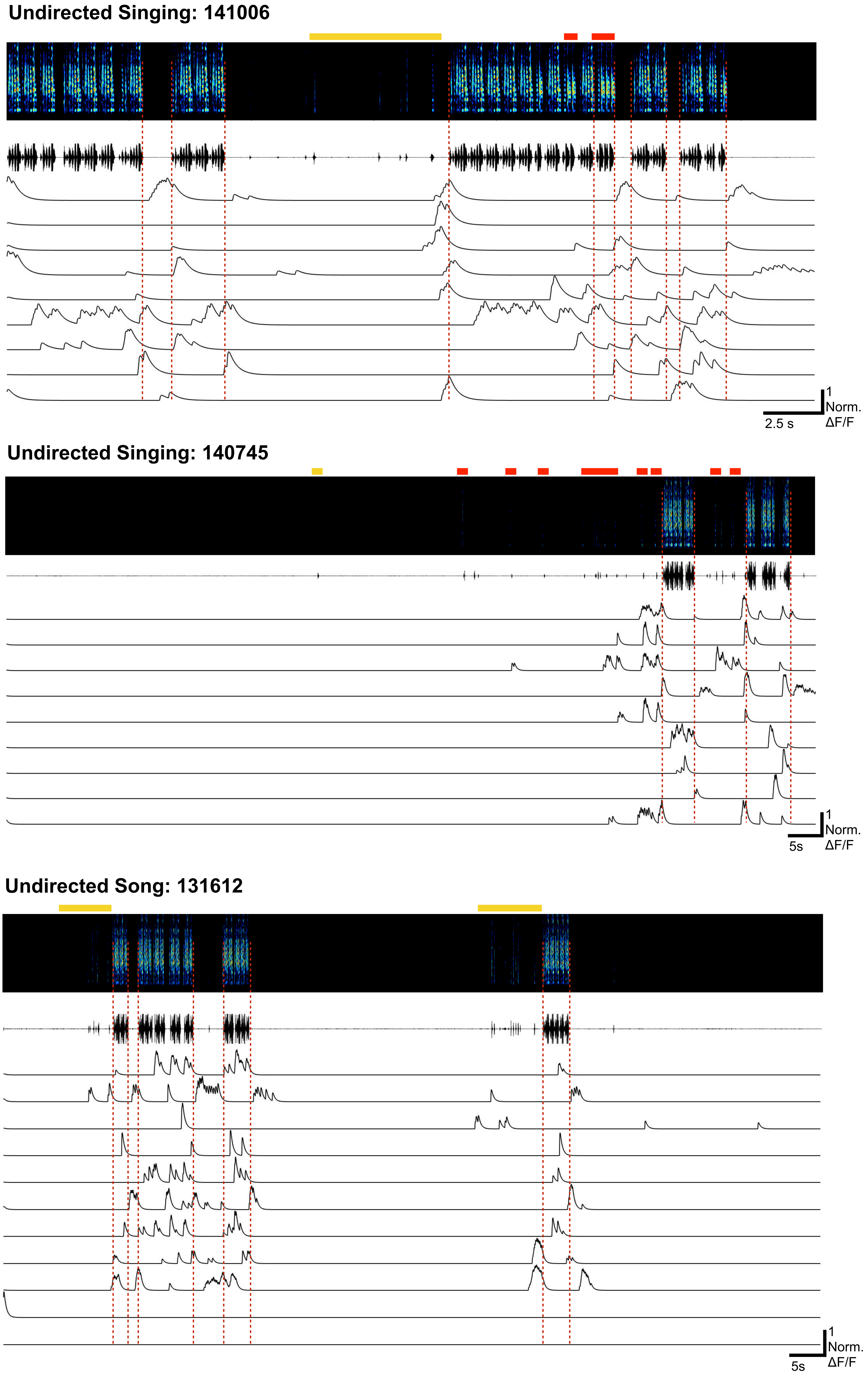


**Figure 6**. Synchronized calcium traces for all available trials for one bird (O248) across directed, undirected, and non-singing behaviors. Bars above spectrograms indicate cage noise (yellow), female calling (blue), and male calling (red). Each trace under the spectrograms correspond to different neurons but they are not necessarily the same neuron across the trials.

### Extended Data Figure 7

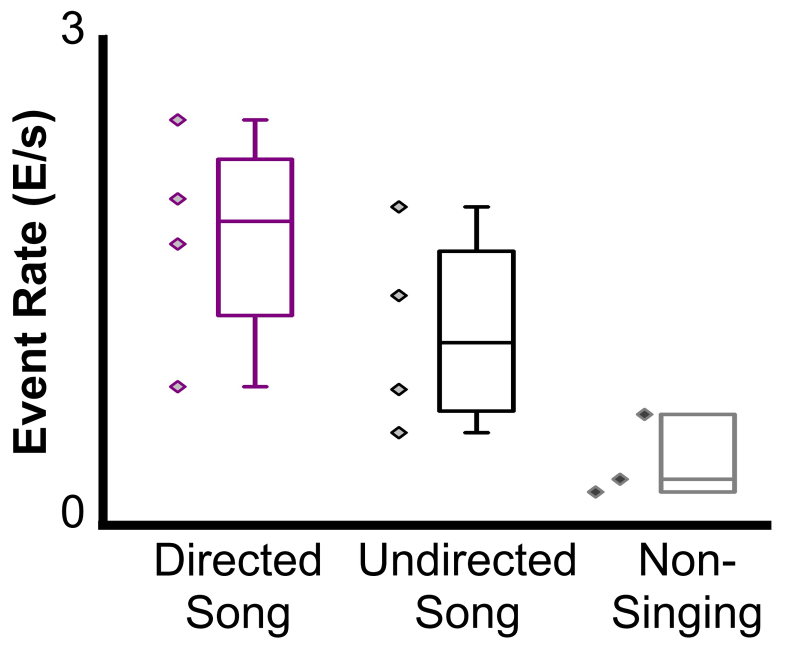


**Figure 7**. a) Event rates for a bird during Directed (561 CEs, 1.67 ± 0.74 std ), Undirected (252 CEs, 1.06 ± 0.66), and Non-Singing (53 CEs, 0.19 ± 0.27) periods of behavior (Directed song is significantly different from non-singing, $F(2,10) = 4.83$, $p < 0.05$). Each marker on the plot corresponds to a different trial. Event rate was calculated as the ratio of the number of events to the duration of the trial.

### Extended Data Figure 8

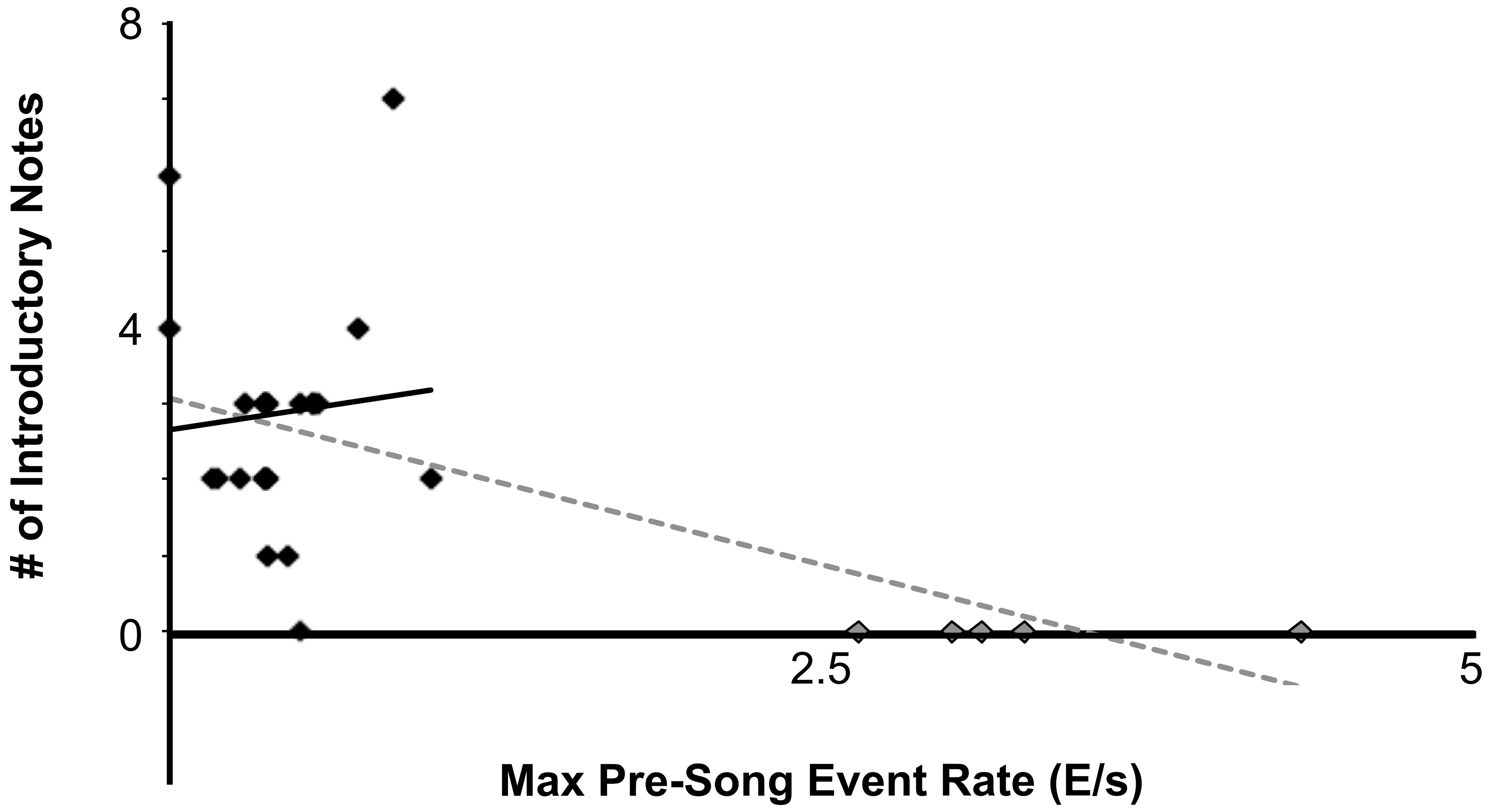


**Figure 8**. Comparison between maximum pre-song event rate to the number of introductory notes prior to motif onset across 5 birds (Mean Event Rate = 0.91 E/s, Mean Introductory notes = ~2). Black trend line excludes grey points (All from one bird that did not have any introductory notes, $r^2 = 0.0078$). Grey dashed trend line includes grey points ($r^2 = 0.34$).

**Extended Data Figure 9**

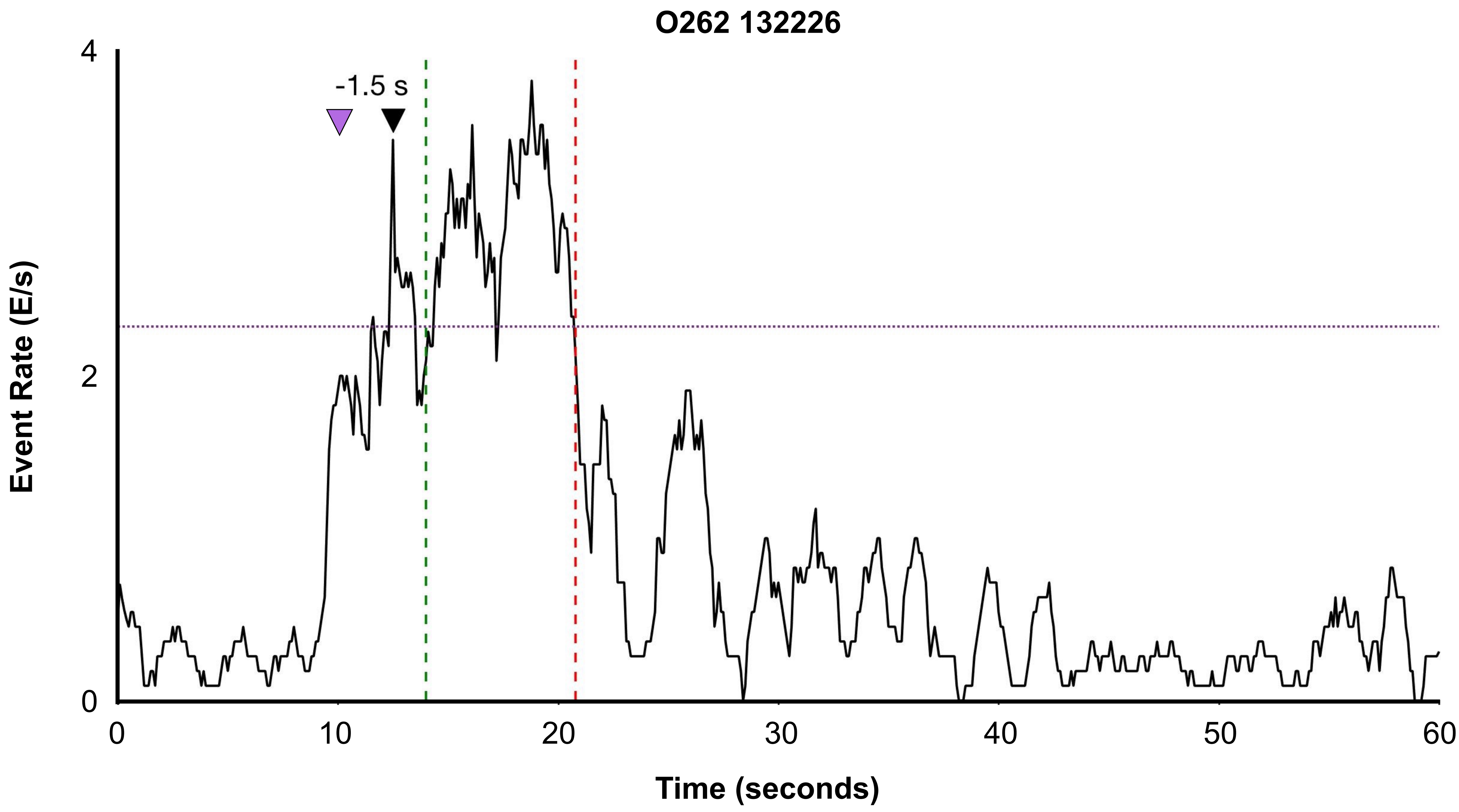


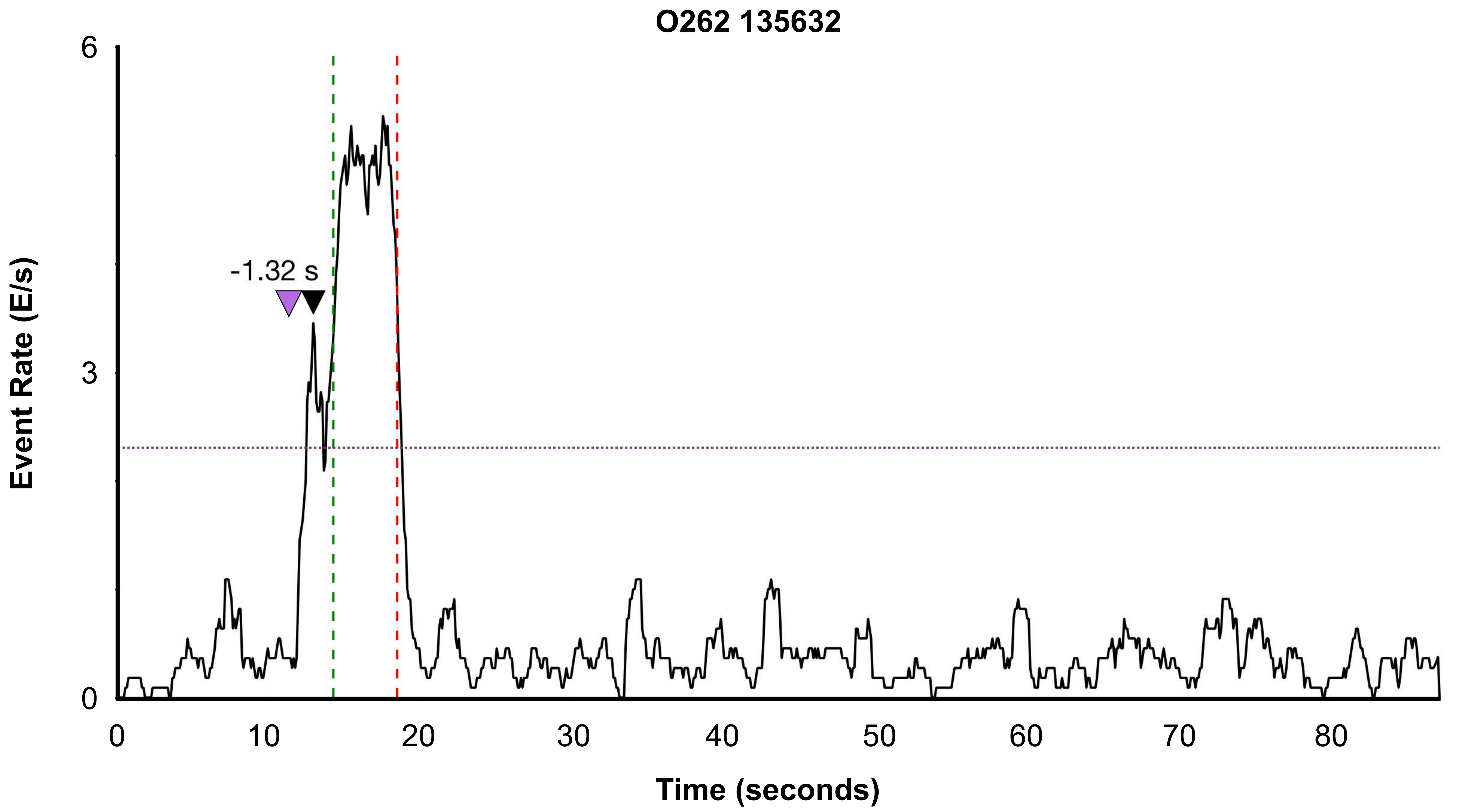

**Extended Data Figure 9**

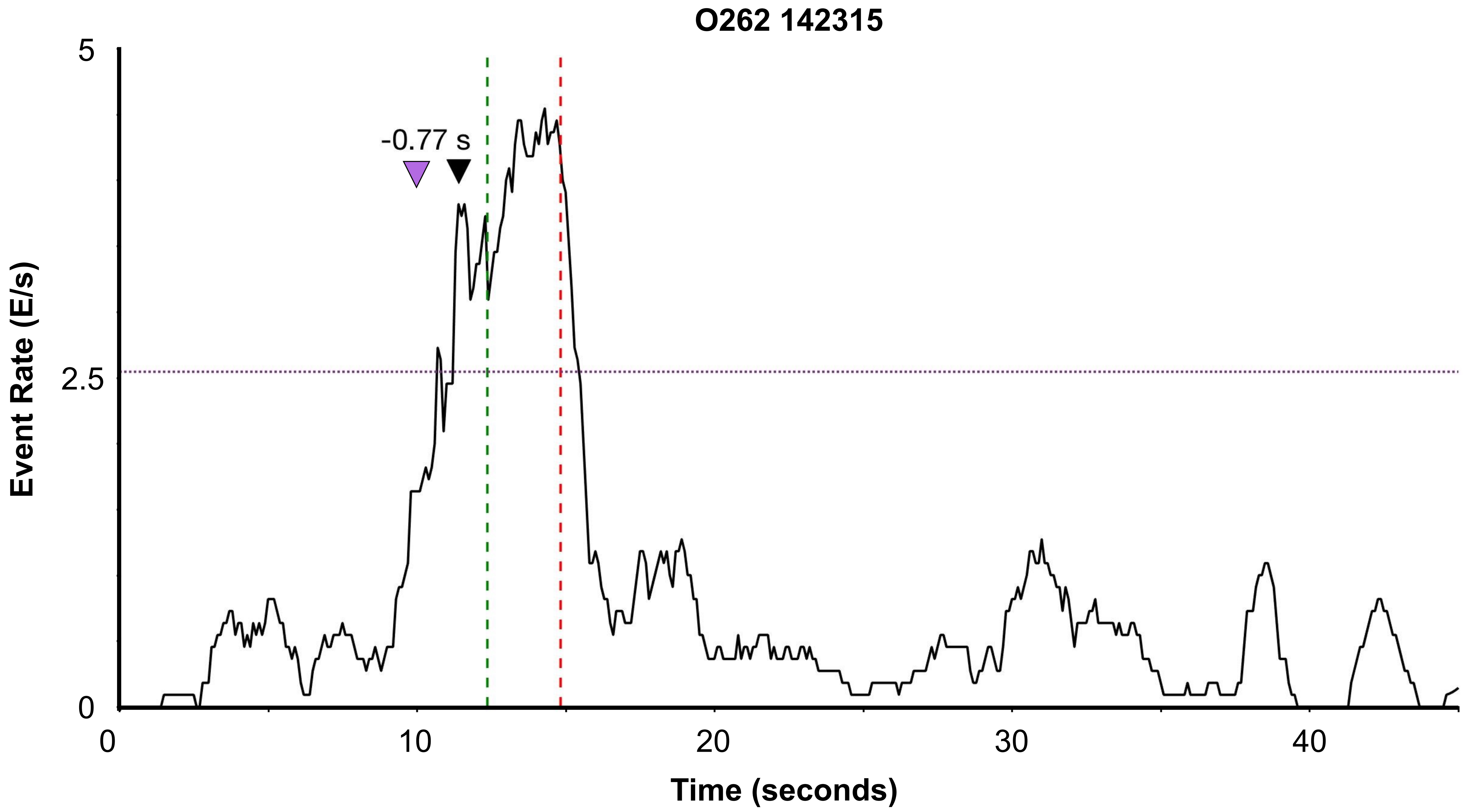


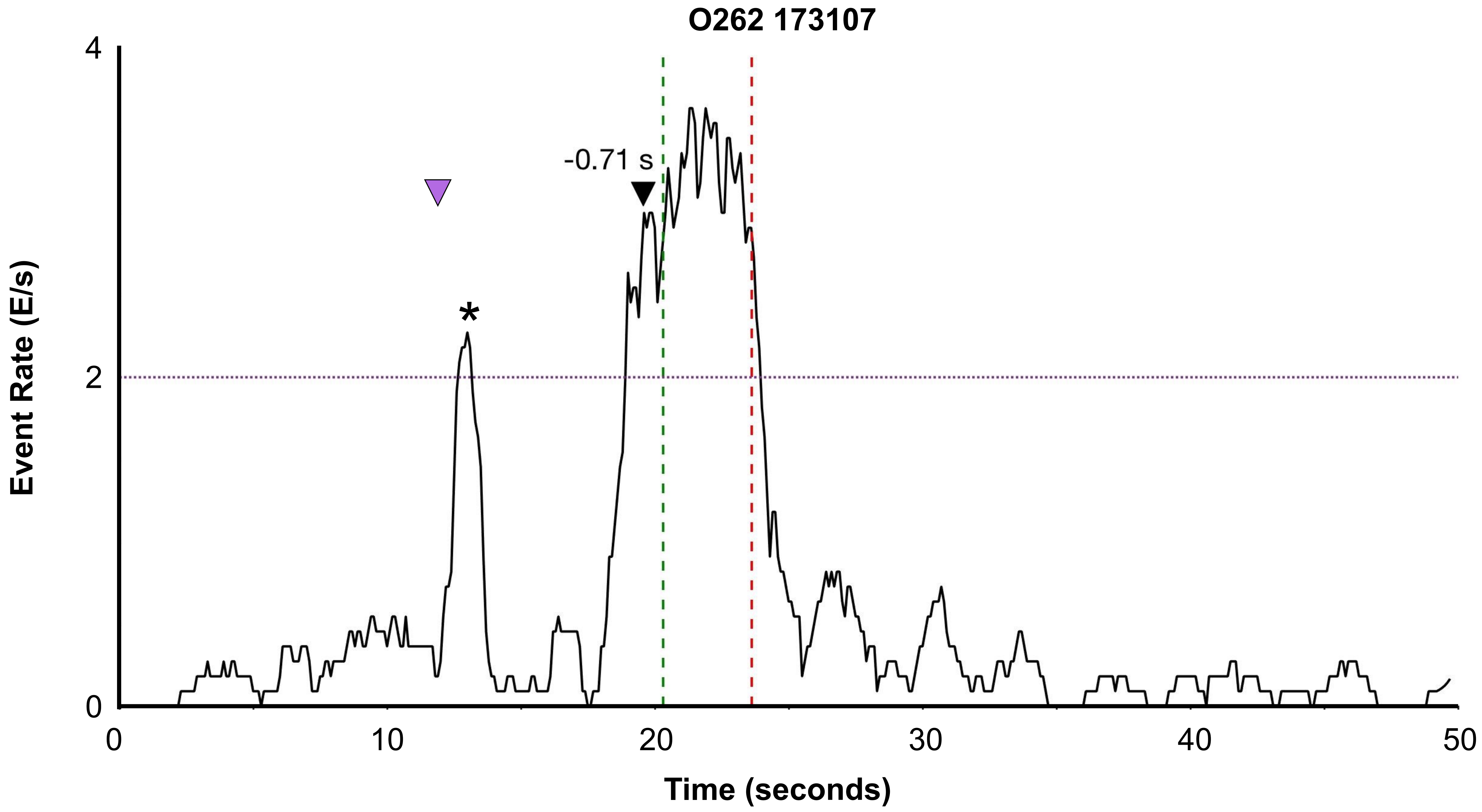

### Extended Data Figure 9

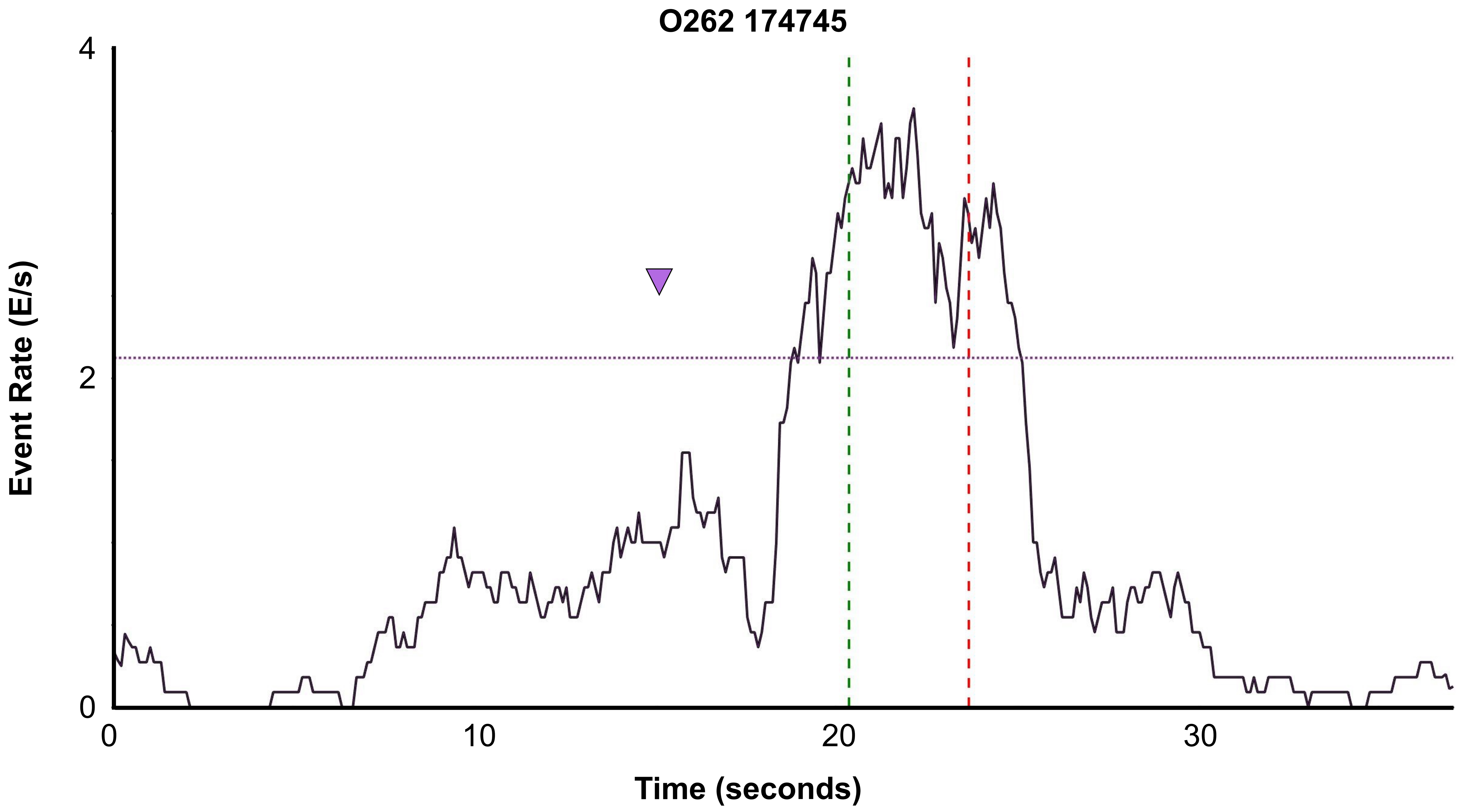


**Figure 9**. Event rate calculated across the full length of 5 trials for 1 bird (O262). Event rate was calculated by first binning calcium onsets in 100ms bins and smoothing with a 1 second moving window. Green dashed line indicates onset of song phrase and red dashed line indicates offset of song phrase. Purple (horizontal) line indicates 2/3s of the maximum value during the pre-song period. Black arrow with time marks maximum peak song event rate. Magenta arrow indicates when female was introduced. Asterisk indicates onset of short call.

### Extended Data Figure 10

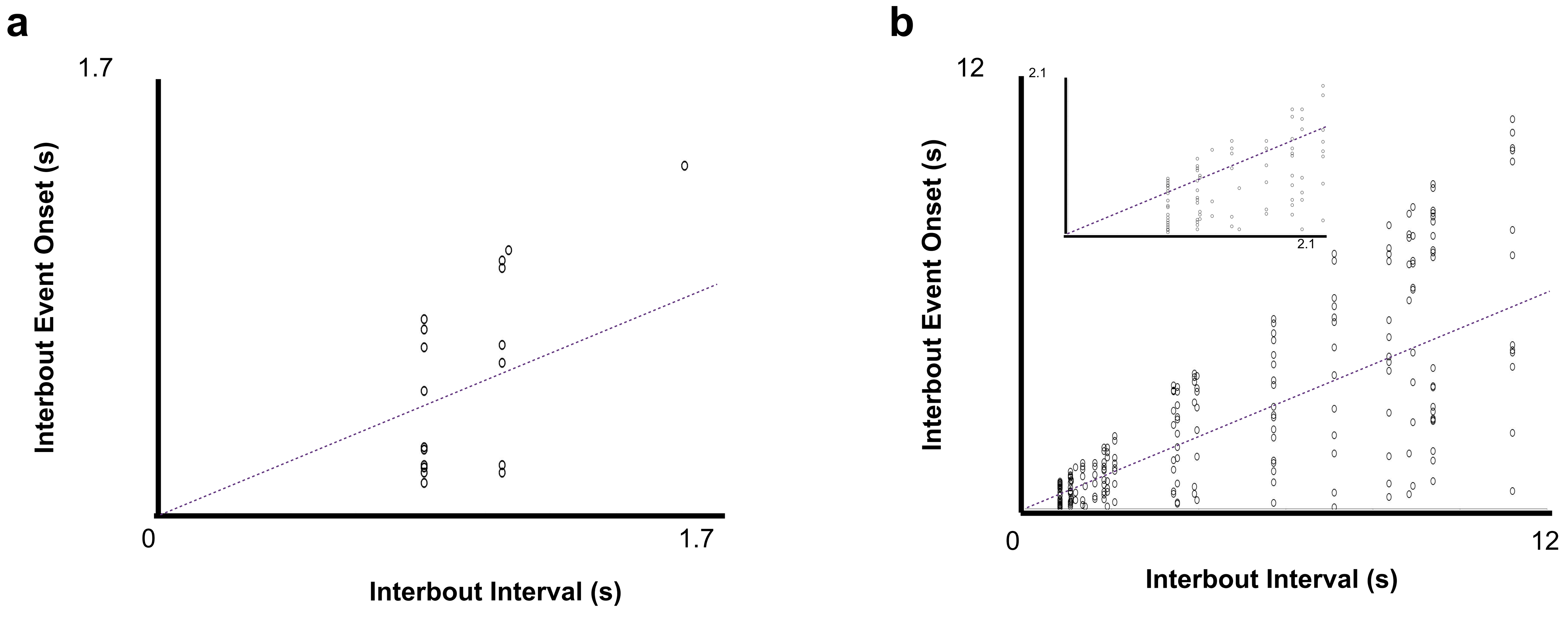


**Figure 10**. a) Inter-bout events for peri-song neurons (N = 27 events, each event was aligned to the end of the previous bout). Dashed line indicates midway point between start of next bout and end of previous bout. These events were excluded from the analysis shown in Figure 3. b) Same as A but for pan-song neurons (N= 231 events). The inset shows a zoomed in portion of events occurring during Interbout Intervals less than 2 seconds. These events were excluded from the analysis shown in Figure 3.

### Extended Data Figure 11

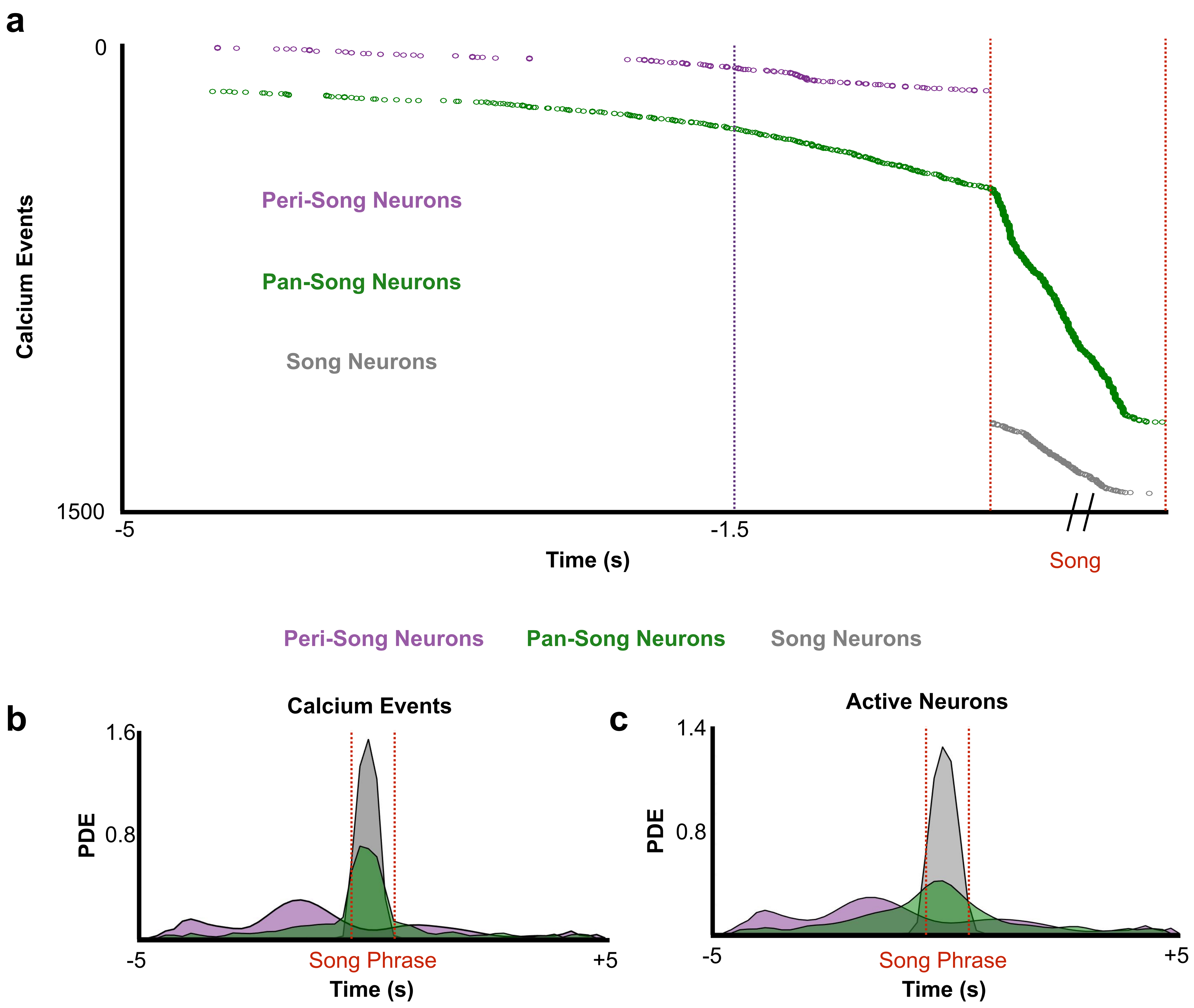


**Figure 11**. **a)** Cumulative calcium event onset times across all birds and all trials. Time corresponding to peak event rate for peri-song neurons is shown (magenta line, 1.5 s). **b)** Probability density estimates of calcium events organized by neuron type. (pan-song neurons: 1,333 CEs, 143 neurons from 6 birds; peri-song neurons: 190 CEs, 41 neurons from 3 birds). **c)** Same as **b** but for active neurons (pan-song neurons: 853 CEs, 143 neurons, 5 birds; peri-song neurons: 169 CEs, 41 neurons, 3 birds).

### Extended Data Figure 12

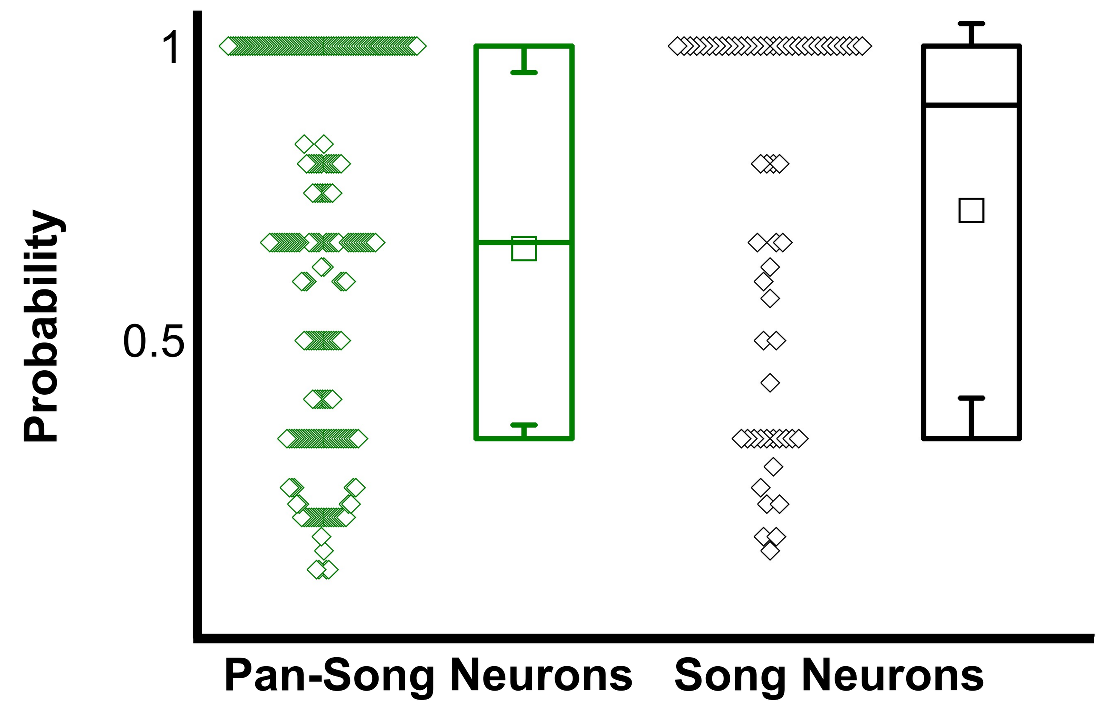


**Figure 12**. Comparison of the probability of pan-song neuron and song neuron events occurring during a motif (Kolmogorov-Smirnov test, n.s., $p > 0.05$; song neurons P(motif) = 0.72 ± 0.32, pan-song neurons P(motif) = 0.66 ± 0.29). Bouts consisting of a minimum of 2 motifs where the neuron was active during at least one of those motifs were used to calculate the probability (29 bouts had greater than 1 motif out of a possible 32 bouts; 28 song neurons; 132 pan-song neurons). Probability was calculated at the bout level for each neuron, probabilities across different bouts were treated as independent variables.
